# The SARS-CoV-2 envelope PDZ Binding Motif acts as a virulence factor disrupting host’s epithelial cell-cell junctions

**DOI:** 10.1101/2024.12.10.627528

**Authors:** Flavio Alvarez, Guilherme Dias De Melo, Florence Larrous, Lauriane Kergoat, Batiste Boeda, Vincent Michel, Danielle Seilhan, Magali Tichit, David Hing, David Hardy, Etienne Kornobis, Hervé Bourhy, Nicolas Wolff, Célia Caillet-Saguy

**Author notes:** These authors contributed equally (Guilherme Dias De Melo, Florence Larrous). These authors contributed equally (Hervé Bourhy, Nicolas Wolff, Célia Caillet-Saguy). Address correspondence to Flavio ALVAREZ.

## Abstract

Severe Acute Respiratory Syndrome Coronavirus 2 (SARS-CoV-2), the virus responsible for the COVID-19 pandemic, has significantly impacted global health, emphasizing the need to understand its pathogenicity and virulence mechanisms. SARS-CoV-2 disrupts the alveolar epithelial barrier and exacerbates airway inflammation, leading to acute respiratory failure, but the molecular details remain unclear. Additionally, SARS-CoV-2 infection causes neurological symptoms, potentially due to its weakly understood ability to cross the blood-brain barrier. The viral multifunctional Envelope (E) protein is crucial for its virulence, playing a key role in virus assembly, budding, and release. The E protein contains a PDZ-binding motif (PBM) that interacts with host PDZ domain-containing proteins, potentially affecting host signaling pathways and contributing to pathogenicity.

This study focuses on the E protein PBM and its role in virulence, disrupting respiratory epithelial barriers and exacerbating airway inflammation. We generated recombinant mutant viruses lacking the PBM and conducted both *in vitro* and *in vivo* experiments to elucidate its impact on viral fitness, pathogenicity, and effects on the epithelial integrity. *In vitro,* the viral mutants showed delayed replication and reduced cytopathic effects. *In vivo,* experiments with hamsters revealed that PBM-deficient viruses caused less weight loss, lower viral loads, and reduced inflammation, indicating decreased pathogenicity. Histological analyses confirmed less airway damage in these hamsters compared to those infected with the wild-type virus. Additionally, PBM-deficient viruses had impaired interactions with tight junction proteins like ZO-1, a PDZ-containing protein, crucial for maintaining epithelial barrier integrity.

Our findings also demonstrate that the PBM does not play a significant role in neuroinvasion during the acute phase of infection, as evidenced by comparable viral RNA loads across brain regions in infected hamsters, regardless of PBM presence. Histopathological and transcriptomic analyses further support this observation, suggesting that the PBM primarily affects specific epithelial barriers. Additionally, RNA-seq analysis on lung and brainstem from infected hamsters reveals that the PBM modulates inflammatory and immune responses, with a stronger impact in lung tissue than in the brainstem. PBM-deficient viruses induce lower levels of inflammation and cytokine expression, suggesting PBM’s specific role in enhancing viral pathogenicity through the activation of pathways such as NF-κB and TNF.

Thus, the E protein PBM plays a critical role in SARS-CoV-2’s fitness, virulence, and pathogenicity, through the disruption of cell junctions and inflammation, underscoring its potential as a target for therapeutic interventions.

## Introduction

Severe Acute Respiratory Syndrome Coronavirus 2 (SARS-CoV-2), the causative agent of the coronavirus disease 2019 (COVID-19) pandemic, has posed a significant threat to global health. This pandemic has highlighted the critical need to understand the molecular mechanisms governing the virus’ pathogenicity and virulence. Among its various pathogenic effects, SARS-CoV-2 has been implicated in disrupting the alveolar epithelial barrier ^1,2^ and exacerbating airway inflammation ^3–5^, which can lead to acute respiratory failure. However, the intricate molecular mechanisms governing these effects remain incompletely understood while they are essential for identifying potential therapeutic targets.

SARS-CoV-2 is an enveloped virus with a positive strand RNA genome encoding for non-structural and structural proteins essential for its replication and pathogenicity. Among the latter, the multifunctional Envelope (E) protein emerges as a critical virulence factor ^6–8^, orchestrating crucial aspects of the viral life cycle such as virus assembly, budding, and release. Studies on both SARS-CoV-1 and SARS-CoV-2 have demonstrated that viruses lacking the E protein exhibit significantly reduced titers, underscoring the protein’s necessity for virus particle formation ^9–11^. While extensive research has been made in understanding the roles of other structural proteins such as the Spike, the Membrane, and the Nucleocapsid proteins, the specific mechanisms and effects of the SARS-CoV-2 E protein on virulence, epithelial junctions, and mucosal inflammation remain less well understood and warrant further investigation ^12,13^. The E protein of SARS-CoV-2 contains a type 2 PDZ-Binding Motif (PBM) at its C-terminus (DLLV_COOH_) with the consensus sequence X-φ-X-φ_COOH_ (where X represents any amino acid and φ a hydrophobic residue), identical to the one of SARS-CoV-1 ^7^. The SARS-CoV-1 and 2 PBM is known to interact with host PDZ domain-containing proteins, modulating host cell signaling pathways and contributing to the virus’s pathogenic profile. Indeed, previous studies have underscored the harmful impact of high viral loads on airway epithelial tight junctions, involving for instance interactions with Proteins Associated with Lin Seven 1 (PALS1) and Syntenin ^7,14,15^.

In addition to airway damage, SARS-CoV-2 infection is characterized by a wide array of acute and chronic neurological symptoms, including headaches, seizures, attention disorders, sleep disorders, short-term memory loss, and neuropsychiatric symptoms such as anxiety and depression ^16–18^. These neurological aspects of the infection are particularly intriguing as the mechanisms of neuroinvasion are not well understood and may involve the crossing of cerebral barriers, such as the blood-brain barrier (BBB) and the blood-cerebrospinal fluid barrier (BCSFB) ^19,20^. These structures, especially the choroid plexus, are rich in PDZ-containing proteins such as ZO-1, which are essential for cellular junctions^21,22^. The E protein of SARS-CoV-2 has been reported to upregulate inflammation pathways in the brain, accompanied by depression-like behaviors and dysosmia when injected directly into the brain ^23^. The disruption of cell junctions remains controversial in the literature ^24,25^, but the PBM of the SARS-CoV-2 E protein might play a pivotal role in this process and raises questions concerning the specificity of epithelial disruptions during infection.

In our previous high-throughput interactomic screening, we identified ten human PDZ-containing proteins that bind to the SARS-CoV-2 E protein PBM ^26^. Many of these proteins, such as ZO-1 (also called TJP1), PARD3, MLLT4 (also called Afadin), LNX2, and MPP5 (also called PALS1), play crucial roles in cellular junctions and polarity. These PDZ-PBM interactions were characterized by structural and functional approaches. These PBM-dependent interactions trigger the colocalization and sequestration of PDZ domains with the full-length E protein in cells, specifically in the Golgi compartment ^27^.

In this study, we aimed to elucidate the specific role of the E protein PBM in SARS-CoV-2 virulence and pathogenicity. We generated and characterized two recombinant viruses lacking the PBM through deletion (E-ΔPBM) or mutation (E-MutPBM). Both *in vitro* and *in vivo* experiments demonstrated that the absence of the PBM significantly impaired viral fitness, resulting in delayed replication and reduced cytopathic effects. In golden hamsters, PBM-deficient viruses led to milder symptoms, less weight loss, and reduced airway inflammation and lesions, as confirmed by histological and transcriptomic analyses. Using human bronchial epithelial models, we found that the E protein’s subcellular localization is independent of the PBM, but the PBM is essential for interacting with the tight junction proteins ZO-1, thereby affecting epithelial barrier integrity during infection. While further research is needed to understand the mechanisms of neuroinvasion and epithelial barrier crossing, our findings suggest that the E protein PBM does not significantly affect neuroinvasion processes. Together, these results emphasize the essential role of the E protein PBM in SARS-CoV-2 fitness, virulence, and pathogenicity, highlighting its potential as a target for therapeutic strategies.

## Results

### The alteration of the E protein PBM significantly affects the SARS-CoV-2 viral fitness

To characterize the role of the E protein PBM in the virulence and pathogenicity of SARS-CoV-2, we produced recombinant viruses lacking the E protein PBM using a reverse genetic method based on yeast recombination as previously described ^28^. We generated a recombinant Wild-Type virus from the original Wuhan strain (rSARS-CoV2-E-WT), a deleted construct missing the four last carboxy-terminal amino acids (rSARS-CoV2-E-ΔPBM), and a mutated construct with a quadruplex of glycine instead of the PBM (rSARS-CoV2-E-MutPBM) **(Figure 1A)**. The corresponding infectious RNAs of the full-length virus genomes were electroporated into Vero-E6 cells and rescue viruses from the culture medium were collected and titrated **(Figure 1B).** The viral sequences were controlled by Next Generation Sequencing (NGS) **(Supplementary Table 1)**. All the mutations close to the PBM and elsewhere in the genome above a frequency of 10% in the viral population were analyzed **(Supplementary Table 1)**. 12% of the viral population exhibit a T2087A amino acid substitution in the G2M domain of NSP3 coding sequence, a position in a domain that is weakly characterized in the literature, nor identified in SARS-CoV-2 variants. However, 3% of the viral population of the rSARS-CoV2-E-ΔPBM carried the P71L substitution, a mutation characteristic of the Beta variant B.1.351 ^29^. This early appearance of the mutation suggests significant mutational pressure on this residue, located just one position upstream of the PBM **(Figure 1A)**. All the following experiments were conducted with the initial viral stock (P0 rescue) in which the genomic sequences were consistent. Interestingly, a marked differential in lysis plaque size was observed between the mutants and the WT viruses in the lysis plaque titration assay **(Figure 1B)**. Indeed, PBM-deleted viruses induced the formation of smaller lysis plaques than those induced by the WT virus. This effect was even more drastic in the presence of the PBM-mutated viruses. To assess the involvement of the PBM in virus fitness *in cellulo*, viral growth kinetics of recombinant viruses lacking the PBM were compared to the ones of the WT viruses in Vero-E6 cells **(Figure 1C)**. We verified beforehand that viral growth curves of the SARS-CoV-2 original virus (Wuhan) **(Figure 1C, Wuhan CoV-2 curve)** and the recombinant rSARS-CoV-2-E-WT **(Figure 1C, E-WT curve)** were similar. We observed replication defects for the E-ΔPBM and E-MutPBM viruses at a MOI (Multiplicity Of Infection) of 0.01 during the early stages of infection (prior to 48 hpi, hours post-infection) with a viral production around 2 logs lower than WT viruses. After 48 hpi, all the viruses reached similar titers. This replication delay was even more drastic for an infection at a MOI of 0.001, where, akin to the lysis plaque assay, the E-ΔPBM mutant appeared to exhibit an intermediate phenotype with a viral growth curve between those of the E-WT and the E-MutPBM viruses. As SARS-CoV-2-induced cytopathic effects damage the cell monolayer, we performed impedance measurements in infected Vero-E6 monolayers using the Maestro Z platform (Axion BioSystems) **(Figure 1D)**. To differentiate levels of virus replication and cytolysis effects, confluent Vero-E6 monolayers in CytoView-Z plates were infected with recombinant viruses at multiple MOIs (0.0001 to 1), and resistance measurements were acquired over 96 hpi. The observed shift in sigmoid infectious dose-response curves for PBM-lacking mutants at 72 hpi **(Figure 1D)** and the differential median time to cell death **(Figure 1E)** agreed with a replication delay and attenuated cytopathic effects observed for the mutants. The E-ΔPBM mutant exhibited again an intermediate phenotype with a resistance dose response located between those of the E-WT and the E-MutPBM viruses. Together, these findings suggest that while the E PBM may not play an essential role in viral replication, its absence leads to a delay in viral growth, consistent with the known functions of the E protein in viral assembly, budding, and release.

**Figure 1.**
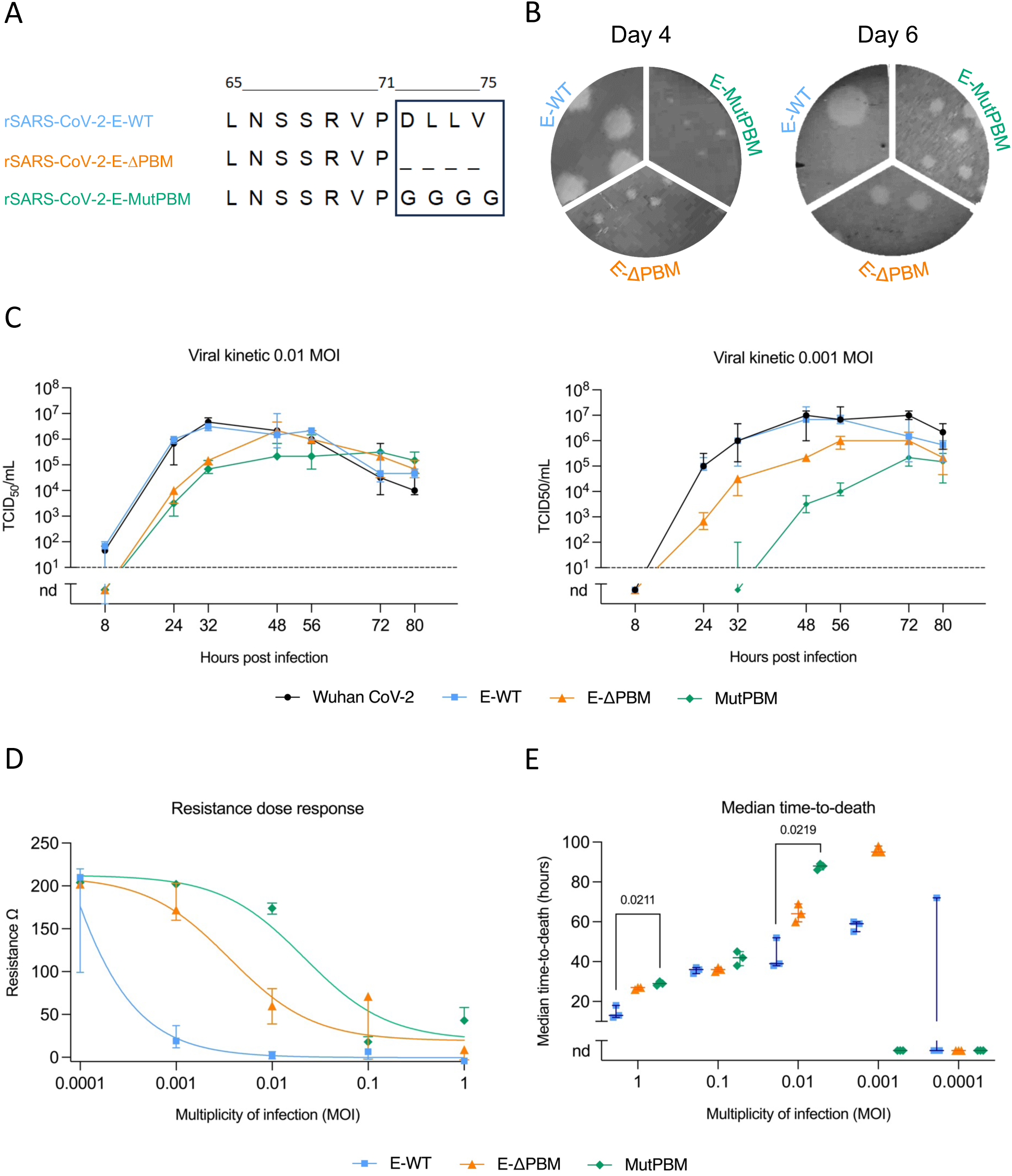
Production and characterization of recombinant SARS-CoV-2 viruses lacking E protein PBM. **A.** Sequences of the Envelope C-terminus end of the recombinant viruses generated by reverse genetics. The box highlights the PBM. rSARS-CoV-2-E-WT, rSARS-CoV-2-E-ΔPBM and rSARS-CoV-2-E-MutPBM correspond to the Wuhan wild-type recombinant virus, and respectively to the viruses lacking the PBM by deletion of the four last residues or by mutations in a quadruplex of glycine. **B.** Plaque morphology of the recombinant viruses. Plaques were developed in Vero-E6 cells on day 4 (left) or 6 (right) after infection. **C.** *In vitro* viral growth curves of the different recombinant SARS-CoV-2 viruses and the original Wuhan SARS-CoV-2 virus (Wuhan CoV-2). Subconfluent monolayers of Vero-E6 cells were infected at an MOI of 0.01 (left) or 0.001 (right). Culture supernatants were collected at several time post infection and titrated by TCID_50_ assay. Dots correspond to means with standard deviations and dashed lines indicate the limit of detection (n=3 independent replicates/time-point, nd: not detected). **D.** Resistance dose response in Vero-E6 cells at 72 hpi. Impedance was measured every minute over the course of 96 hours in wells that were mock infected or infected with recombinant SARS-CoV-2 viruses in 10-fold dilutions ranging from an MOI of 1 to 0.0001. Sigmoids curves were generated using AxIS Z software. Dots indicate the median and vertical lines indicate the interquartile range (n=3). **E.** Median time-to-death calculations based on raw resistance data for each MOI. Horizontal lines indicate median with the interquartile range (n=3). Kruskal-Wallis test followed by the Dunn’s multiple comparisons test (the adjusted p value is indicated when significant).

### The PBM does not impact E protein localization in cells and air-liquid reconstructed human bronchial epithelium

To investigate the localization of the E protein and the impact of the PBM, we infected A549 ACE2 TMPRSS2 cells and MucilAir™ model, an *in vitro* reconstructed human pseudostratified bronchiolar epithelium mimicking the air-liquid interface in the lungs **(Figure 2A)**, with E-WT, E-ΔPBM, and E-MutPBM viruses. The kinetics of viral replication in the apical compartment of the MucilAir™ model showed a replication defect for the E-ΔPBM and E-MutPBM viruses during the early stage of infection, with a viral production 3-4 logs lower than the one of E-WT virus **(Figure 2B)**. From day 4 to day 7 post-infection, the viral loads of the mutant viruses remain stable and never reach the levels observed with the E-WT virus. The E-ΔPBM mutant exhibits an intermediate viral replication kinetic, in line with those observed in Vero-E6 cells. Additionally, a lower viral load was detected for E-WT virus in the basolateral compartment **(Figure 2C)**, indicating that SARS-CoV-2 particles are predominantly released from the apical side of the epithelium. Interestingly, the E-MutPBM virus was not detected in the basolateral compartment, and the E-ΔPBM virus again exhibits an intermediate phenotype with 1 to 2 logs lower than the E-WT virus at 4 dpi. Immunofluorescence analysis was performed on A549 ACE2 TMPRSS cells and on the MucilAir™ model at 2 and 4 dpi respectively, after infection with E-WT, E-ΔPBM, and E-MutPBM viruses **(Figure 2D and E; Supplementary Figure 1).** E protein co-stained with GM130, a marker for the Golgi apparatus. A cytoplasmic vesicular distribution of the E protein is also observed **(Figure 2D and E; Supplementary Figure 1)**, consistent with the secretory pathway of SARS-CoV-2 viral particles, mainly described in the literature through peroxisomes and lysosomes. Regardless of the virus, E proteins are detected on the apical surface of ciliated cells **(Figure 2D)**, distinctly identified from basal cells via actin staining (phalloidin) **(Figure 2E)**. Within the MucilAir™ model, E protein also exhibits intracellular localization, displaying a Golgi-like and vesicular distribution **(Figure 2D and E)**. Thus, the different subcellular localization observed with both E-WT and E mutant viruses indicates that E localization is independent of the PBM in infected cells or in the MucilAir™ model.

**Figure 2.**
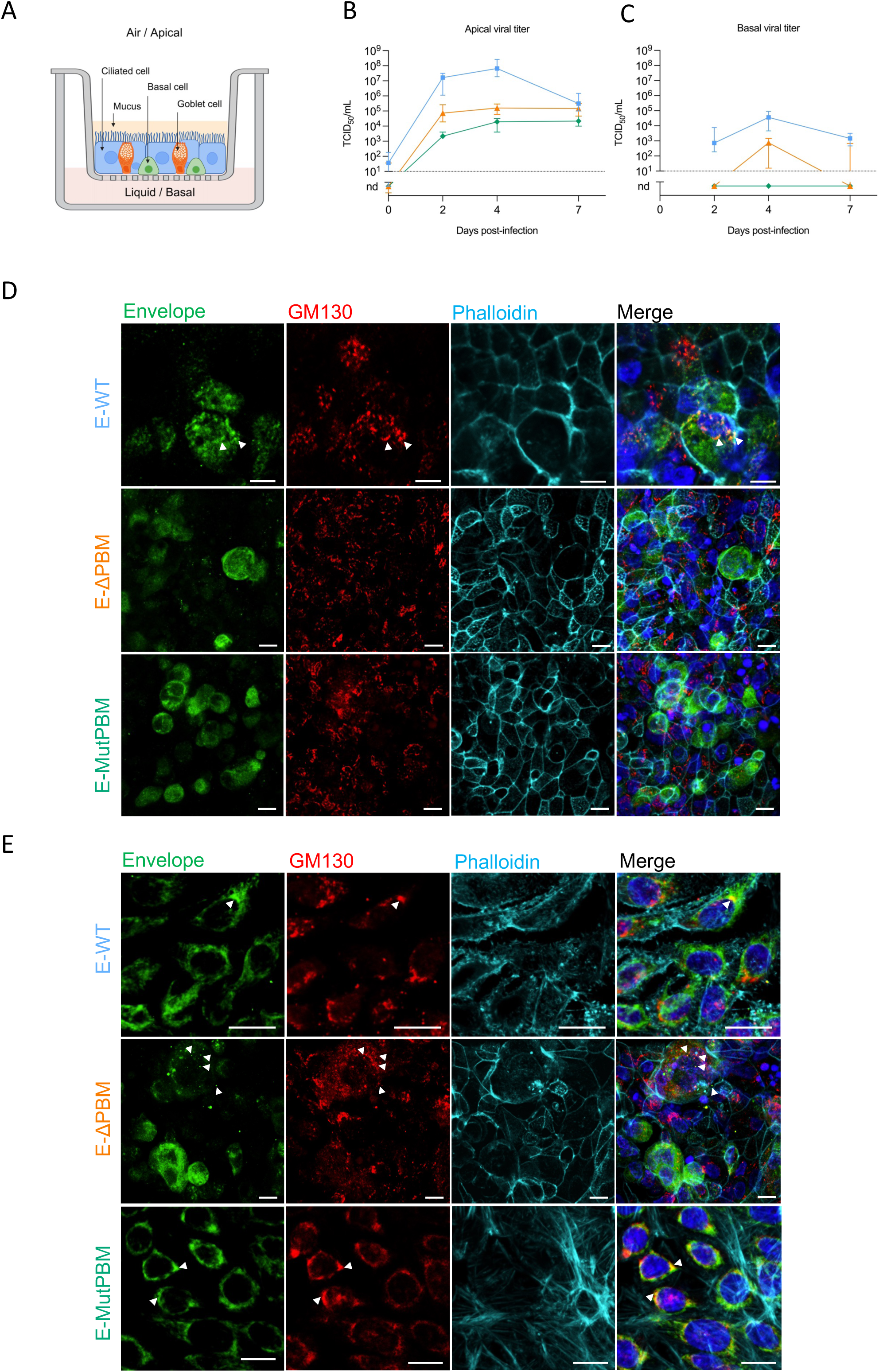
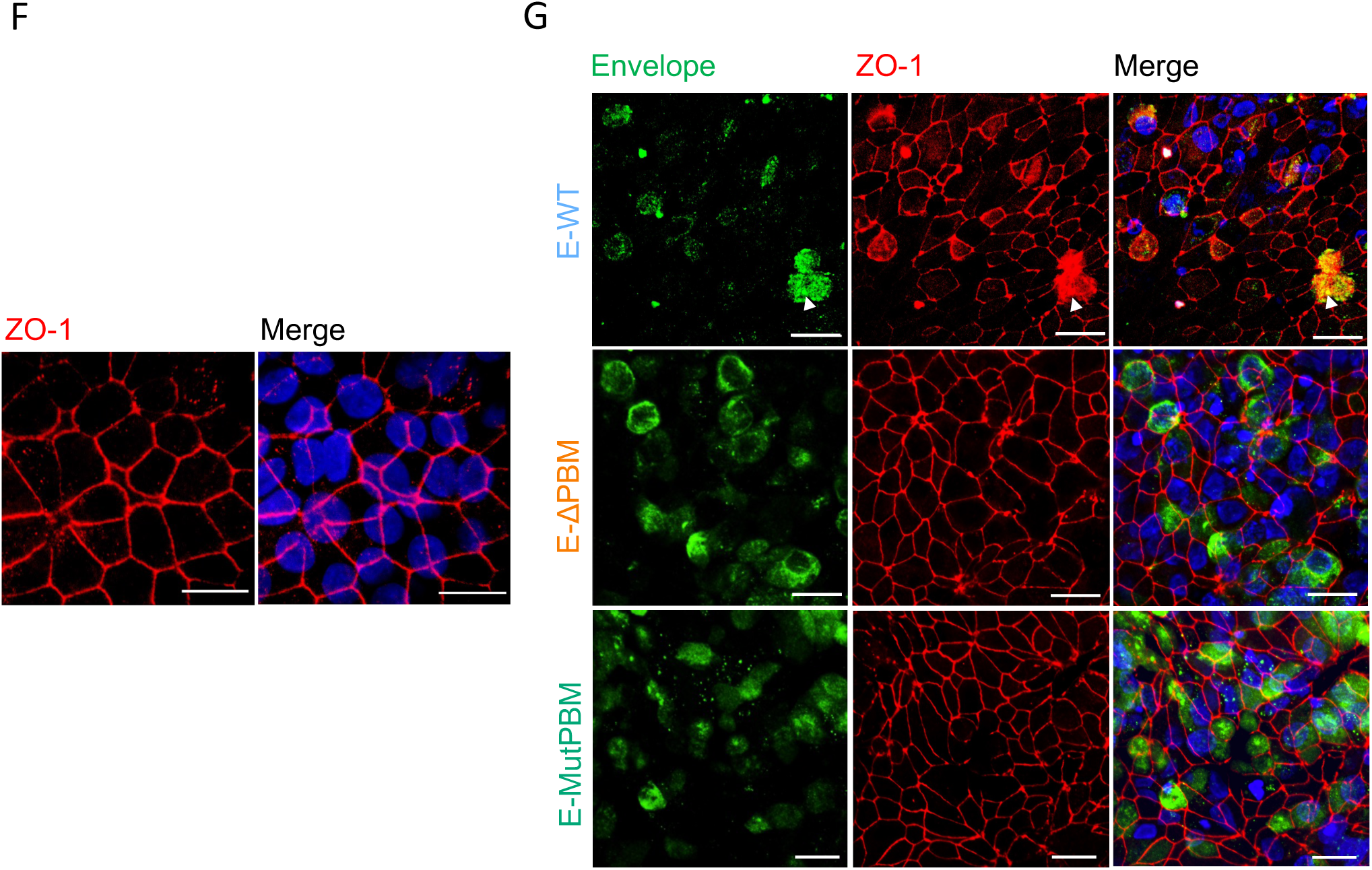
Subcellular localization of the Envelope protein and subversion of ZO-1 protein during SARS-CoV-2 infection in a Lung-on-Chip model. **A.** Schematic view of the MucilAir reconstructed human bronchial epithelium. The model exhibits an air-liquid interface with different cell types (ciliated, goblet, and basal cells). **B.C.** Kinetics of viral replication of the recombinant viruses in apical (B) and basal (C) compartment measured by TCID50 assay at 2, 4 and 7 dpi. Day 0 titration time point corresponds to the 3^rd^ wash supernatant viral quantification. Dots correspond to medians with interquartile range and dashed lines indicate the limit of detection (n=3 independent replicates/time-point, nd: not detected). **D.E.** Immunofluorescence assay showing E protein localization within apical ciliated cells **(D)** and basal cells **(E)** in the MucilAir model. Staining was performed at 4 dpi on a 250 000 pfu infected epithelium with recombinant E-WT (top panel), E-ΔPBM (middle panel), or E-MutPBM (bottom panel) (scale bar = 10 µm). Images are reconstituted in 3D from Z stack acquisitions. Hoechst: nuclei (blue); GM130: Golgi apparatus (red); Envelope: recombinant SARS-CoV-2 envelope protein (green); Phalloidin: actin (cyan). Arrowheads indicate E-Golgi colocalizations. **F.G.** Visualization of ZO-1 protein network correlated with E protein localization in the MucilAir model using immunofluorescence staining. Staining was performed on either non-infected epithelium **(F)** or on epithelium fixed at 4 days post-infection (dpi) with 250,000 pfu of E-WT, E-ΔPBM or E-MutPBM recombinant viruses **(G)**. Arrowheads indicate areas of E-ZO-1 colocalization. Images are reconstituted in 3D from Z stack acquisitions (scale bar = 20 µm). Hoechst: nuclei (blue); ZO-1: zonula occludens 1 (red); Envelope: recombinant SARS-CoV-2 envelope protein (green).

### The SARS-CoV-2 E protein interacts with ZO-1 in a PBM-dependent manner, leading to cell-cell junction alterations during infection

We then investigated the localization of ZO-1 and its interaction with the E protein upon infection with E-WT virus in A549 ACE2 TMPRSS cells **(Supplementary Figure 2A-E)** and the MucilAir™ model **(Figure 2F and G; Supplementary Figure 2F and G)**. Immunofluorescence assays of mock-infected A549 ACE2 TMPRSS cells and MucilAir™ model showed the characteristic ZO-1 staining intact pattern at cell junctions **(Figure 2F; Supplementary Figure 2A)**. In A549 ACE2 TMPRSS cells, when the E-WT protein distribution was observed at the Golgi apparatus, the ZO-1 pattern appears largely unimpaired. However, in regions where E protein presents a sporadic cytoplasmic distribution, meaning an advanced step in the viral cycle within the cell, ZO-1 staining is disrupted supporting a epithelial reorganization **(Supplementary Figure 2B)**. A similar behavior was observed in the MucilAir™ model **(Figure 2G; Supplementary Figure 2F)**. Colocalizations between E and ZO-1 proteins were observed in various subcellular locations, related to Golgi organelle, cytoplasmic vesicles, and cell surface **(Figure 2G; Supplementary Figure 2C and F)**. Interestingly, cells infected with viruses lacking PBM exhibited significantly reduced disruption of ZO-1 **(Figure 2G; Supplementary Figure 2D, E and G).** To confirm the interaction between E and ZO-1 proteins in a PBM-dependent manner, *pull down* assays were performed. GFP-ZO-1, or GFP alone as control cell lysate, were incubated with lysates from cells infected with E-WT or E-MutPBM viruses. The E-WT protein is strongly precipitated with GFP-ZO-1 **(Supplementary Figure 2H)** while the E protein with a mutated PBM is not retained by GFP-ZO-1. These results indicate that E protein produced by viruses in cells interacts with full-length ZO-1 in a PBM-dependent manner. Together, these data support the role of the E protein/ZO-1 interaction in the tight junction disturbance and the impairment of epithelial barrier integrity during SARS-CoV-2 infection.

### PBM-defective viruses infection mitigates the severity of clinical disease associated with SARS-CoV-2 infection in hamsters

After validating the role of the E protein in SARS-CoV-2 pathogenicity *in vitro*, we further investigated the involvement of E PBM in SARS-CoV-2 pathogenicity *in vivo*, by intranasally inoculating male golden hamsters with 6.10^4^ PFU of the three recombinant viruses (E-WT, E-ΔPBM and E-MutPBM). Daily monitoring of body weights **(Figure 3A; Supplementary Figure 3A)** and clinical signs **(Figure 3B; Supplementary Figure 3B)** allowed assessment of disease progression. While E-WT infected hamsters present a progressive weight loss and the onset of clinical symptoms, the animals infected with the PBM-lacking viruses displayed no or minor weight loss and clinical signs, comparable to non-infected hamsters. Lung-to-body weight ratio indicates a significant attenuation of lung inflammation in PBM-deficient infected animals, closely matching the profile observed in non-infected hamsters **(Figure 3C; Supplementary Figure 3C)**. In all these parameters, E-ΔPBM mutant displays again an intermediate phenotype. As anosmia constitutes one of the clinical signs of COVID-19, olfactory performance was assessed at 3 dpi. Interestingly, half of the animals infected with E-WT viruses exhibited anosmia (2/4, *p value =0.0007* from Chi square statistical test), whereas those infected with the two mutant viruses fully conserved their olfactory capacity **(Figure 3D; Supplementary Figure 3D)**. Altogether, these results demonstrate that removal of the E protein PBM drastically decreases pathogenicity *in vivo* highlighting that a functional PBM in E protein is essential for SARS-CoV-2 virulence.

**Figure 3.**
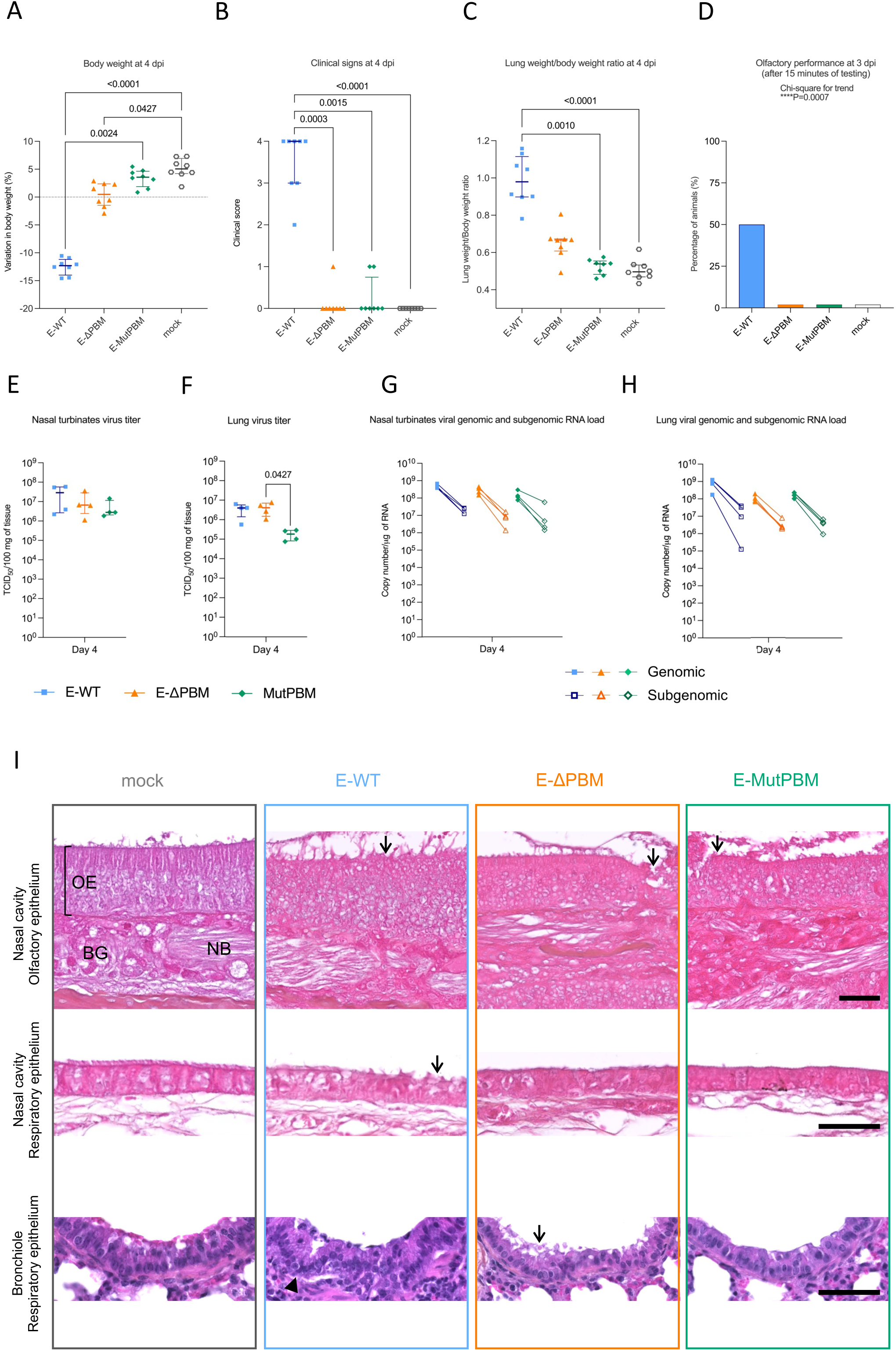
Clinical profile and viral metrics of hamsters infected with recombinant wild-type SARS-CoV-2 (E-WT) and PBM lacking viruses (E-ΔPBM and E-MutPBM). **A**. Body weight at four days post-infection (4 dpi). **B.** Clinical score at 4 dpi. The clinical score is based on a cumulative 0–4 scale: ruffled fur; slow movements; apathy; and absence of exploration activity. **C.** Lung-to-body weight ratio measured at 4 dpi. Horizontal lines indicate median and the interquartile range (n=8/group). Kruskal-Wallis test followed by the Dunn’s multiple comparisons test (the adjusted p value is indicated when significant). **D.** Olfactory performance loss measured at 3 days post-infection (dpi). The olfaction test is based on the hidden (buried) food finding test. Bars represent the percentage of anosmic animals (n= 4/group). Chi-square test for trend. **E.F.** Infectious viral titers in nasal turbinates **(E)** and lung **(F)** at 4 days post-infection (dpi) expressed as TCID_50_ per 100 mg of tissue. Horizontal lines indicate median and the interquartile range (n=4/group). Kruskal-Wallis test followed by the Dunn’s multiple comparisons test (the adjusted p value is indicated when significant). **G.H.** Viral genomic and subgenomic RNA load detected in nasal turbinates **(G)** and lung **(H)** at 4 dpi. Horizontal lines indicate median and the interquartile range. Lines connect symbols from the same animals (n=4/group). Kruskal-Wallis test followed by the Dunn’s multiple comparisons test (the adjusted p value is indicated when significant). (n=4/group). **I.** Histopathological study of airway epithelia from hamsters infected with wild-type SARS-CoV-2 recombinant virus (E-WT) and PBM-lacking viruses (E-ΔPBM and E-MutPBM) at 4 dpi. Representative images of Hematoxylin and Eosin (H&E) staining, showing the olfactory epithelium (top panels) and respiratory epithelium (middle panels) in the nasal cavity from full sagittal sections of hamster heads, as well as the bronchiolar epithelium (bottom panels) from whole lung sections. Arrows indicate epithelium’s damages and arrowhead correspond to inflammation. Scale bar = 50 µm; n = 4 per group. Abbreviations: OE = olfactory epithelium; BG = Bowman’s gland; NB = nerve bundles (*filia olfactoria*).

To assess the effect of the E protein PBM in virus growth *in vivo*, viral loads were quantified by titration and RT-qPCR viral RNA quantification at 1, 2 and 4 dpi in the upper and lower airways (nasal turbinates and lung) **(Figure 3E-H; Supplementary Figure 3E-H)**. While substantial viral charges were found in both airways’ organs for E PBM-deficient viruses (10^5^-10^7^ TCID_50_/100 mg of tissue), E-WT virus replicates at higher level (10^7–8^ TCID_50_/100 mg of tissue). Lower quantities of viruses were recorded for E-ΔPBM and E-MutPBM viruses at the early-stage post infection mainly at 1 and 2 dpi. This tendency was more pronounced in lung than in nasal turbinates, consistent with the direction of the viral spread into the respiratory tract. An intermediate profile was again observed for the E-ΔPBM virus located between those of the E-WT and E-MutPBM viruses **(Supplementary Figure 3E-H)**. Genomic viral RNA in lung and nasal turbinates of all animals were detected with loads at 10^8–9^ copy number/μg of RNA associated with subgenomic RNA loads at 10^5–9^ copy number/μg of RNA **(Figure 3G and H; Supplementary Figure 3G and H)**, in agreement with the replicative capacity of all viruses *in vivo*.

To further explore the role of the E protein PBM beyond the respiratory tract, we assessed its involvement in viral infection within the brain, a site known to be affected by SARS-CoV-2, comparing its effects with those observed in the airways. Within the brain, titration in the different brain region at 4 dpi (olfactory bulb, cortex, brainstem and cerebellum) highlights the detection of infectious viral particles only in the olfactory bulb of animals infected with E-WT virus **(Supplementary Figure 4A)**. In contrast to the titration results, no differences were observed between the viruses when detecting genomic RNA in the olfactory bulb and cortex of the animals with high genomic RNA loads at 10^7^ and 10^8^ copy number/μg of RNA respectively. More variability between replicates was observed in the brainstem and the cerebellum, in which several RNA loads could not be quantified. Except one positive hamster in the brainstem for the E-WT animal infected group, no subgenomic RNA was detected in the different parts of the brain (**Supplementary Figure 4B)**. Thus, the absence of a functional PBM delayed the viral growth in the airway tract, while its implication in the neuroinvasion seems minor with progressive infection through the olfactory bulb, cortex, brainstem, and cerebellum.

### Damages to upper and lower airways epithelia are significantly reduced with PBM lacking viruses compared to E-WT virus

Histological analysis was performed on nasal cavity and lungs sections from infected hamsters with E-WT, E-ΔPBM, and E-MutPBM viruses at 4 dpi **(Figure 3I and Supplementary Figure 5)**. Examination of E-WT infected nasal epithelia sections reveals disorganization with important inflammation and desquamation. The bronchial epithelium shows substantial damage characterized by subsidence and desquamation **(Figure 3I)**. Across all levels of airway epithelia, E-ΔPBM-infected samples exhibit moderate pathogenic signs, whereas E-MutPBM samples show only minor signs, comparable to those observed in non-infected animals **(Figure 3I)**. These histological findings suggest that the observed attenuation in viruses lacking E PBM is correlated with a decrease in the airway’s pathology. Examination of lung sections from hamsters infected with E-WT exhibits significant edema, congestion, thickening of alveolar walls, and severe perivascular and peribronchiolar mononuclear cells infiltrates **(Figure 3I; Supplementary Figure 5A)**. Additionally, viral antigen detection by immunostaining with SARS-CoV-2 nucleocapsid antibody reveals widespread distribution of the virus within the organ **(Supplementary Figure 5B)**. In contrast, hamsters infected with viruses lacking E PBM show intermediate phenotypes between the mock-infected hamsters and those infected with E-WT, with significantly less damage and lung edema. The E-ΔPBM mutant displays an intermediate phenotype between those of E-WT and E-MutPBM **(Figure 3I; Supplementary Figure 5A)**. SARS-CoV-2 nucleocapsid staining demonstrates limited viral propagation in the organ for E-ΔPBM and E-MutPBM compared to E-WT **(Supplementary Figure 5B)**.

To focus on epithelial structures potentially disrupted by SARS-CoV-2, Hematoxylin-Eosin staining was performed on whole brain sections of hamsters at 4 dpi. We focused our investigation on brainstem and choroid plexus structures. No significant morphological changes were observed during infection, and the choroid plexus appeared similar across all groups (**Supplementary Figure 4C**). Immunohistochemical (IHC) staining for the glial fibrillary acidic protein (GFAP), an astrocyte marker, indicated an absence of inflammation when comparing mock-infected and E-WT-infected animals (**Supplementary Figure 4C**). To assess potential blood-brain barrier leakage, fibrinogen IHC staining was performed, revealing no differences at this time point of the infection (4 dpi) **(Supplementary Figure 4C)**. Altogether, these findings suggest that SARS-CoV-2 neuroinvasion is highly restricted, with no evidence of blood-brain barrier leakage, microscopic brain damage, or astrocyte activation.

### The PBM drives a distinct transcriptomic signature, primarily modulating inflammatory signaling proteins and affecting adherens junction components

To investigate the PBM’s impact of the PBM on SARS-CoV-2 virulence, we conducted a comparative RNA-seq transcriptomic analysis during the peak of infection, at 4 dpi, in the lungs and brainstem of hamsters infected with the E recombinant viruses **(Figure 4; Supplementary Figure 6)**. Principal Component Analysis (PCA) and Venn diagrams **(Figures 4A and 4B)** revealed that the transcriptomic responses (with an absolute fold change > 2) were significantly more pronounced in the lungs compared to the brainstem, consistent with the SARS-CoV-2’s respiratory tropism. In the lungs, over 1,000 genes were differentially expressed compared to non-infected hamsters, while only a few dozen genes were impacted in the brainstem **(Figure 4B)**.

**Figure 4.**
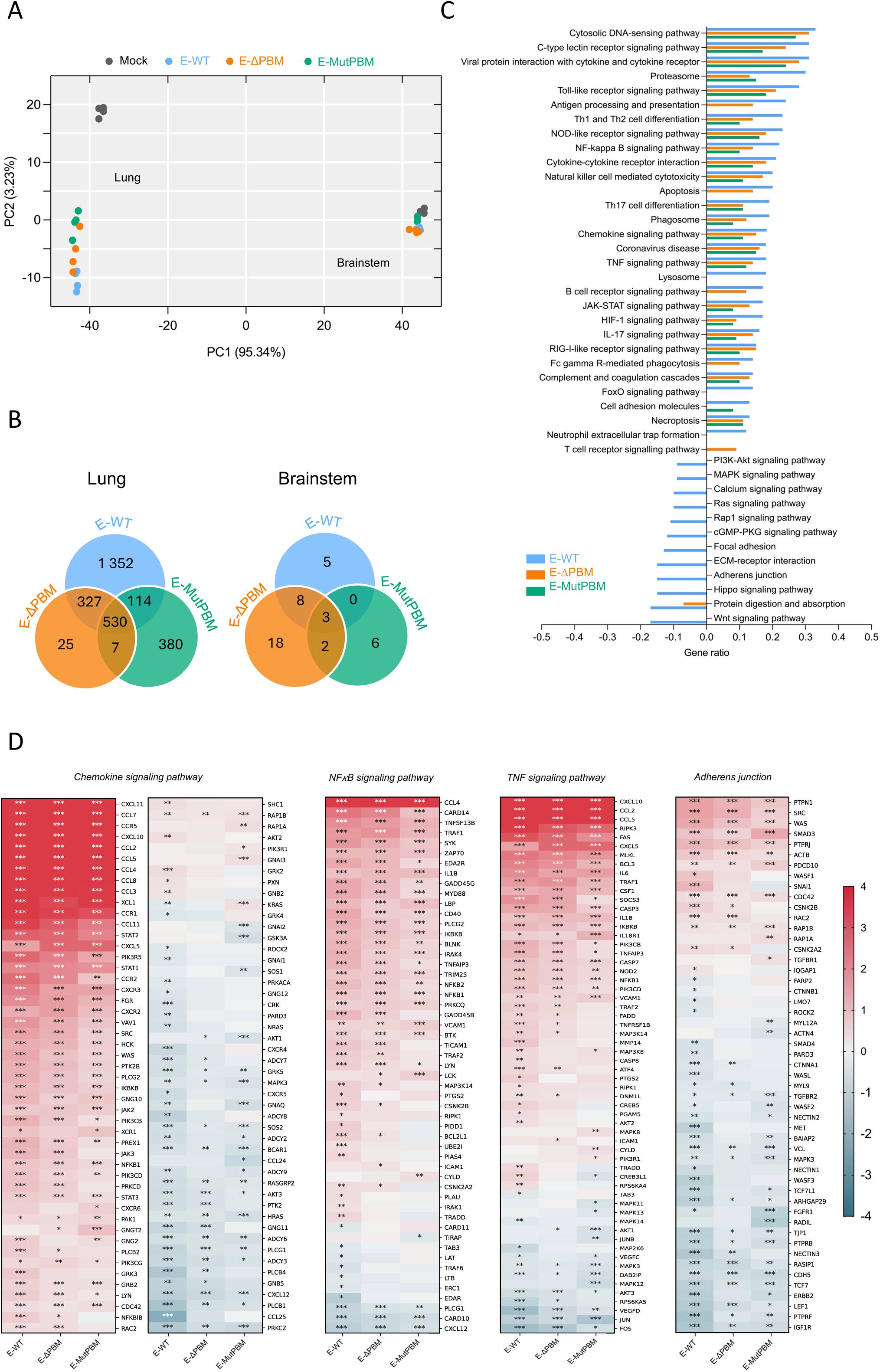
Transcriptomic analysis of lung and brainstem responses to PBM-lacking viruses’ infection in hamsters. **A.** Principal component analysis (PCA) of the general transcriptome characteristics. Transcriptomic analysis was conducted on lung and brainstem tissues from hamsters that were either non-infected (mock) or infected with E-WT, E-ΔPBM, and E-MutPBM viruses at 4 days post-infection (dpi). The first principal component (PC1) explained 95.34% of the total variance in the dataset, while the second principal component (PC2) accounted for an additional 3.23%. **B.** Venn diagram showing the number of differentially expressed genes. The Venn diagram displays the number of differentially expressed genes, with numbers indicating both unique and shared DEGs when comparing the various recombinant-infected lung (left) and brainstem (right) tissues to non-infected controls (with an absolute fold change > 2). **C.** KEGG enrichment analysis in lung tissue. The histogram shows the relevant KEGG pathways that are significantly up- or down-regulated in recombinant virus-infected lung tissue compared to non-infected controls, ranked by gene ratio. The gene ratio (x-axis) represents the proportion of significant genes relative to the total number of genes within each pathway. **D.** Heatmaps of differentially expressed genes in key KEGG Pathways in lung tissue. Heatmaps display the expression levels of all differentially expressed genes in relevant KEGG pathways in the lungs of hamsters infected with E-WT, E-ΔPBM, and E-MutPBM, compared to mock-infected controls at 4 days post-infection (dpi) with an absolute fold change > 2. Asterisks (*) indicate genes with a Benjamini–Hochberg-adjusted p-value < 0.05 in comparisons between each recombinant virus group and the mock-infected group. The color gradient represents the log₂ fold change in gene expression between infected and control animals, with PDZ-containing genes (TJP1 and ERBB2) highlighted in bold.

Notably, the expression of 530 genes in the lung was consistently altered across all three recombinant viruses compared to non-infected animals, reflecting a core SARS-CoV-2 infection signature that is independent of PBM alterations **(Figure 4B)**. This indicates that some pathways are commonly activated during infection, regardless of PBM presence. The different profiles observed in various infection parameters upon infection with the three recombinant viruses were reflected in the transcriptomic data. Indeed, only 32 genes differed between E-WT and E-ΔPBM, while 387 genes were distinct between E-WT and E-MutPBM **(Figure 4B)**. According to the KEGG analysis, several key inflammatory and immune pathways were consistently disrupted across all the three viruses, including "cytokine-cytokine receptor interaction", as well as the "TNF", "TLR", "NF-κB ", and "chemokine” signaling pathways **(Figure 4C and D)**. In addition, pathways related to cell death mechanisms, including “apoptosis”, “necroptosis”, “proteasome”, and “phagosome formation” were also dysregulated. In PBM-deficient viruses, although the same inflammatory pathways were enriched, they were less intensely modulated compared to the E-WT virus in terms of gene number and foldchange. This matches the reduction in tissue inflammation observed through histopathological analysis **(Figure 3I, Supplementary Figure 5A)**. To validate these findings, we conducted RT-qPCR at 1, 2, and 4 dpi in the upper and lower airways on common pro-inflammatory cytokines (*MX2*, *CXCL10*, *IL-6*, *IFN-B* and *TNF-α*), known to be upregulated during SARS-CoV-2 infection **(Supplementary Figure 6A)**. By day 4, RT-qPCR showed that cytokine expression levels in mutant-infected animals were comparable to those of E-WT-infected animals. However, at day 1, viruses lacking the E protein PBM showed significantly lower expression (up to 20-fold) compared to E-WT-infected animals. This reduction in cytokine expression was most pronounced in the lungs compared to nasal turbinates, notably for *MX2* and *CXCL10*, which showed the highest overexpression at day 1. This decrease in inflammatory cytokine levels reflects delayed immune responses in agreement with the reduced viral fitness and pathogenicity of PBM-deficient viruses *in vitro* and *in vivo* **(Supplementary Figure 6A)**.

Moreover, several crucial signaling pathways, such as "RAS," "MAPK," "WNT," and "PI3K-AKT," were downregulated in E-WT infected animals but unchanged in those infected with PBM-deficient E mutants **(Figure 4C)**. Pathways specifically associated with PBM effects, such as "lysosome function", "neutrophil extracellular trap formation", "antigen processing and presentation", and “apoptosis”, were also identified. In the lungs of E-WT-infected animals, adherens junction proteins, notably those with PDZ domains such as *ZO-1 (TJP1)* and *ERBB2*, were downregulated compared to mock-infected controls.

As initial differences in viral load were observed in the brainstem, a key regulator of vital functions, we centered our transcriptomic analysis on this region. The results revealed no major pathway dysregulation, with only a few genes showing differential expression upon the infection **(Figure 4B)**. This finding aligns with the relatively limited impact of SARS-CoV-2, and specifically the PBM in the brainstem, indicating a finely tuned regulatory response in this region. Nonetheless, an inflammatory response was observed across all three groups, characterized by upregulation of key inflammatory genes, including *ISG15*, *IRF*7, *MX1* and *MX2* **(Supplementary Figure 6B and C)**. Differences in inflammatory response between E-WT and PBM-deficient viruses in the brainstem were highlighted as well as several genes such as *FOS*, *KCNV1*, *PLPPR4*, and *EGR3* **(Supplementary Figure 6B and C)**. Notably, several PDZ-related proteins were identified, including *CNKSR2*, *SGK1* and *ERBIN*.

Overall, our findings demonstrate a marked difference in the transcriptional and inflammatory response between the lungs and the brainstem. The lungs showed extensive gene expression changes, while the brainstem’s response is more limited, indicating a finely tuned immune regulation in this region.

## Discussion

In this study, we provided insights into the critical role played by the envelope protein’s PBM in the virulence and pathogenicity of SARS-CoV-2, spanning from molecular characterization to *in vivo* infection in the hamster model. Using reverse genetics, we produced recombinant viruses expressing E protein with either a deleted E-ΔPBM or a mutated E-MutPBM. Titration and viral growth kinetics showed a delay in replication, underscoring the involvement of the PBM in viral fitness, consistent with the role attributed to this protein-protein interaction motif in the literature^30^. Indeed, the PBM of the SARS-CoV-2 E protein plays a role in viral replication and virulence by interacting with host PDZ-containing proteins, inducing an excessive proinflammatory response, and contributing to alveolar edema and acute respiratory distress syndrome.

The pathogenicity is marked by a disruption at the epithelial level of the respiratory tract ^1,31–33^. Airway barriers are characterized by a complex interplay of tight junctions, mucociliary clearance mechanisms, and immune surveillance. Their disturbance highlights a critical aspect of viral pathogenesis: viruses often target epithelium and cellular junctions and polarity components to hijack cellular machinery ^34–36^. Respiratory virus infections are known to dysregulate tight junction protein, compromising barrier function, and facilitating pathogen invasion into the subepithelial space. PDZ domain-containing scaffold proteins are key components of paracellular impermeability and innate immunity by linking transmembrane proteins of epithelial junctions to the cell cytoskeleton ^34,37,38^. The E protein PBM of SARS-CoV-1 has been identified to target the PDZ domain of PALS-1, leading to disruption of cellular structures ^14^. We previously identified PDZ-containing proteins, partners of the PBM involved in cell junction and polarity and characterized these interactions at structural and functional levels ^27^. ZO-1, a protein observed to be mislocalized and disrupted during SARS-CoV-2 infection ^1,24,39–41^, emerges as a key partner for the E PBM.

Before investigating the disturbance of PDZ-containing proteins, we focused our study on the cellular localization of the E protein. Immunostaining in infected A549 ACE2 TMPRSS2 lung epithelial cells and infected Lung-on-Chip model (MucilAir) revealed that the E protein localization depends on an upstream Golgi targeted sequence in the C-terminal tail, unaffected by the PBM ^42,43^. As reported in the literature for SARS-CoV-1 and 2, regardless of whether the viral construct includes the PBM or not, the E protein accumulates in the Golgi apparatus but also exhibits a vesicular cytoplasmic distribution. This generic term might encompass endosomes, lysosomes, peroxisomes, and other vesicular organelles involved in the exocytosis pathway and viral egress ^11,44–47^. Moreover, studies have reported Golgi enlargement in response to the infection, indicating structural changes that may facilitate viral assembly and release ^46,48^.

Focusing on the cell junction disturbance, we previously demonstrated that the E protein PBM interacts with the PDZ2 domain of ZO-1 [26] and that transfected E protein, when expressed at the Golgi apparatus, is able to recruit PDZ2 and full-length ZO-1 to this organelle [27,32]. During infection, ZO-1 staining revealed mislocalization and degradation in A549 cells and the MucilAir model. This phenomenon is directly associated with the PBM, as reported by immunofluorescence colocalization and co-immunoprecipitation experiments. Interestingly, this perturbation depends on the localization of the E protein. No disturbance is observed when the E protein is localized at the Golgi apparatus, suggesting that viral entry is not a critical step for epithelial disruption. However, when the E protein is engaged into the exocytosis pathway and exhibits vesicular localization associated with endosomal and lysosomal organelles, ZO-1 appears disrupted. These findings indicate that the disruption of cellular structures by the E protein is not solely dependent on the PBM but also on the protein’s subcellular localization. It also supports the conclusions drawn from studies on PALS-1 during SARS-CoV-1 infection, where the E protein might impact the trafficking of cell junction components to the membrane, leading to leakage between adjacent epithelial cells, disruption of the epithelial barrier, and ultimately promoting viral spread ^14^. In the context of cells infected with PBM mutant viruses, the disruption of ZO-1 appears significantly diminished. However, its preservation is not complete, as the perturbation of epithelial integrity results from multiple mechanisms involving several viral proteins. While the E PBM is widely recognized as a factor of pathogenicity, the Orf3a and Nucleocapsid proteins also possess a PBM, interacting with specific PDZ domain-containing proteins, with a limited literature ^26,49–51^. Interestingly, our previous *in vitro* high-throughput screening of the Orf3a PBM against the full human PDZome showed that ZO-1 is also a potential interactant ^26^. Deletion of the SARS-CoV-1 Orf3a PBM has minimal impact on replication in infected cells and only a modest effect on weight loss or mortality in mice ^50^. While its role in viral pathogenesis remains unclear, the Orf3a PBM may partially compensate for a PBM-deficient E protein.

A striking attenuation of pathogenicity was observed *in vivo* when hamsters were infected with viruses lacking the E PBM, as previously reported for SARS-CoV-1 and 2 in mice ^7,8,30^. In our study, the animals infected with PBM-deficient viruses exhibited no weight loss and no clinical signs of the disease. Quantification of viral infectious particles revealed substantial viral loads in the airways for PBM-deficient viruses, though at lower levels compared to E-WT infection at the early time points of the infection. This phenomenon was more pronounced in the lower airways, consistent with the virus’s pathway in the respiratory tract and propagation defects observed for the PBM-deficient viruses. Interestingly, the viruses expressing deleted or mutated PBM exhibited different behaviors following various infection parameters. Indeed, the PBM-mutated virus showed a stronger attenuation of phenotypes compared to the PBM-deleted virus. These results suggest a differential host response to the distinct envelope C-terminus sequences resulting from these modifications. Several hypotheses can be formulated concerning this effect. The deletion induces a change in E protein size that might impact various mechanisms in which E is involved. From this perspective, the literature highlights several crucial motifs within the cytoplasmic domain of the E protein involved in protein and lipid interactions, folding, and post-translational modifications ^52–55^. Moreover, this deletion could create a potential alternative internal PBM. Indeed, the deletion provides the sequence -NLNS<u>SRV</u>P_COOH_ with a potential internal type I PBM (-X-S/T-X-ϕ) instead of type II PBM (-X-ϕ-X-ϕ_COOH_) found in the E-WT (-NLNSSRVPD<u>LLV</u>_COOH_). This new PBM might interact with a subset of PDZ-containing proteins targeted by E-WT, as it has been demonstrated that different PBM types in coronaviruses can have overlapping targets ^26^. It is also possible that the new PBM recruits other PDZ-containing proteins, partially compensating for the loss of the original interactions. Another hypothesis might be that the mutation and/or deletion of the PBM in viruses differentially alter the RNA structure in a different way in a region that plays a role in viral translation and/or replication. Further investigation is required to fully understand these changes and their implications.

In addition, our study focused on examining the role of the PBM in anosmia and neuroinvasion. The brain barriers such as the choroid plexus is reported to be highly enriched in PDZ-containing proteins ^21,22^, which tightly regulate the central nervous system (CNS) homeostasis as well as protect the CNS from toxins, inflammation, injury, and pathogens ^56^. This also prompts a broader consideration of epithelial barriers, raising the question of whether the effect of the PBM is specifically linked to the disruption of certain tissues. Unlike the animals infected with the E-WT virus, the hamsters infected with viruses lacking PBM retained their olfactory performance. Moreover, the genomic viral RNA quantification revealed similar viral loads regardless of the virus. These data support the notion that anosmia and neuroinvasion are independent phenomena during infection ^28^. Indeed, anosmia is predominantly associated with the effects of the infection on the olfactory epithelium, where disorganization, inflammation, apoptosis, and damages to the ciliary layer occur ^57,58^.

In this perspective, we focused our analysis on the impact of the PBM on airways lesions. Notable histological phenotypes associated with the infection and the respiratory failure were observed. The hamster lungs infected with E-MutPBM present a dramatic reduction in edema and inflammation compared to animals infected with E-WT, and E-ΔPBM virus again displays an intermediate phenotype exhibiting moderate inflammation. Altogether these data support the literature on the involvement of the PBM and more largely the E protein in the inflammatory process characterizing the COVID-19 ^59–62^. Histopathological imaging of the upper airways, showing a drastic reduction in lesions, is consistent with the preservation of olfactory capabilities in animals infected with the PBM-deficient viruses. Although the olfactory bulb is a common structure in the SARS-CoV-2 cerebral infectious process, the literature also reports infection through the blood brain barriers, including the choroid plexus ^20,63–66^. At 4 dpi, no morphological changes at the macroscopic scale were recorded in the histopathological analysis, nor was there any evidence of astrocyte activation or barrier leakage.

While COVID-19 infection is known to induce a severe inflammatory response, particularly in the early stages, which can lead to respiratory failure and death ^4,67,68^, our transcriptomic analysis further supports that the E PBM plays a critical role in amplifying this response. Indeed, in PBM-deficient cases, inflammatory pathways were enriched in our RNAseq but with notably less intensity in terms of gene number and foldchange. A significant decrease in the expression of genes involved in inflammatory pathways was observed, supporting the role of the PBM in driving the unrestrained immune response typically triggered during SARS-CoV-2 infection ^5,69,70^. These data demonstrate that the exacerbated host innate immune response was impaired and delayed in the absence of PBM, aligning with the viral replication defect observed in airway organs. These findings may explain the attenuated pathogenicity observed for these viruses. The downregulation of key signaling pathways, including “WNT”, “MAPK”, and “PI3K-AKT”, observed exclusively in E-WT-infected lungs suggests a PBM-specific role in modulating these critical cellular processes. This finding warrants further investigation to elucidate potential mechanistic connections, as PBMs in other viruses are known to influence similar pathways ^71–74^. Additionally, these findings correlate with the observed downregulation of adhesion junction proteins, including PDZ-containing proteins such as ZO-1, suggesting that the PBM may impact epithelial integrity not only at the proteomic level but also through gene expression regulation. However, transcriptomic changes in the brainstem, were minimal between PBM-deficient and E-WT virus infections, with the identification of 3 PDZ-related proteins: *CNKSR2*, which features both a PBM and a PDZ domain ^75^, as well as *SGK1*, recognized for its interaction with the PDZ-containing NHERF family ^76^ previously reported as a partner of the E PBM ^26^, and ERBIN, a basolateral epithelial protein ^77^ related to Ras-Raf and NF-kB signaling pathways ^78–81^. Although the presence of PDZ-related proteins was notable, further investigations are needed to fully understand the molecular mechanisms through which the E protein PBM modulates brain function and the pathological consequences.

Thus, the E protein serves as a virulence factor that influences viral fitness, spread, and pathogenicity *in vivo*. Altogether, these observations underscore the critical role of the SARS-CoV-2 E protein in disrupting epithelial barrier function, highlighting the significant and innovative therapeutic potential of targeting the E protein’s PBM and its interactions with PDZ proteins.

### Ethics

All animal experiments were performed according to the French legislation and in compliance with the European Communities Council Directives (2010/63/UE, French Law 2013–118, February 6, 2013) and according to the regulations of Institut Pasteur Animal Care Committees. The Animal Experimentation Ethics Committee (CETEA 89) of the Institut Pasteur approved this study (200023; APAFIS#25326-2020050617114340 v2) before experiments were initiated. Hamsters were housed by groups of 4 animals in isolators and manipulated in class III safety cabinets in the Pasteur Institute animal facilities accredited by the French Ministry of Agriculture for performing experiments on live rodents. All animals were handled in strict accordance with good animal practice.

### Generation of Wuhan SARS-CoV-2 recombinant viruses

The recombinant viral genomes were constructed as previously described by *de Melo et al, 2023*. 11 overlapping fragments around 3kb each and two fragments encoding for yeast-specific selection genes (His3 and Leu2) were recombined in the yeast centromere plasmid pRS416 to generate the complete genome of the recombinant virus. The T7 promotor was placed just before the 5’UTR sequence and a unique restriction site EAG1 was placed just after the poly(A) tail. Viral fragments were obtained either by RT-PCR on RNA extracted from Vero-E6 cells infected by Wuhan SARS-CoV-2 using the SuperScript IV VILO Master Mix (11756050, *Thermo Fisher Scientific*) according to the manufacturer protocol, or using synthetic genes (*GeneArt*). All PCR amplifications were performed using Phusion™ High-Fidelity DNA Polymerase (F530, *Thermofisher*). To produce recombinant viruses deleted or mutated in the PDZ Binding Motif (PBM) of the Envelope protein **(Figure 1)**, the PCR product of the fragment 9 that include E protein sequence was cloned in Topo-TA vector, using TOPO™ TA Cloning™kit (K465001, *Invitrogen*) according to the manufacturer protocol. Deletion of the 4 last carboxy-terminal amino acids in rSARS-CoV2-E-ΔPBM (LNSSRVP) or mutation in rSARS-CoV2-E-MutPBM (LNSRVPGGGG) **(Figure 1A)** were introduced by Site-Directed Mutagenesis following Site-Directed Mutagenesis Phusion Kit manufacturer protocol (F541, *Thermo Fisher Scientific*). TOPO-TA-F9 plasmid was used as a template and mutagenesis PCR were performed with specific primers: E_ΔPBM_FOR 5ʹ-TAAACGAACTAAATATTATATTAGTTTTTCTGTTTGG-3’ and E_ΔPBM_REV 5ʹ-AGGAACTCTAGAAGAATTCAGA-3’ primers pair was employed for PBM deletion and E_MutPBM_FOR 5ʹ-TTCTAGAGTTCCTGGTGGTGGAGGTTAAACGAAC-3’ and E_MutPBM_REV 5ʹ-GAATTCAGATTTTTAACACGAGAGTAAACG-3’ primers pair for the PBM mutation. Recombination of 11 overlapping viral DNA fragments and 3 yeast specific fragments (PRS416, Leucine-2 and Histidine-3) was performed into *S. cerevisiae* BY4740 strand cultured for three days at 30 °C. For each recombination, subcultured clones were checked by Multiplex PCR Kit to control the presence of the different fragments (data not shown) (206143, *QIAGEN*) according to the manufacturer protocol, using specific subsets of pair primer pairs as previously described (*Dias de Melo et al, 2022)*. To perform purification of a large amount of recombinant plasmid, the yeast colony of interest was cultured overnight in 200 mL of SD-His-Ura-Leu-medium at 30 °C under agitation (200 rpm). DNA extraction was processed using the NucleoBond Xtra Midi Plus kit (740422, *Macherey-Nagel*). Manufacturer protocol was followed with a beforehand lysing step in which harvested cells were incubated in 16 mL RES-Buffer complemented with 1.6 mg Zymolyase® 100-T (120493-1, *AmsBio*) and 160 μl of β-mercapthoethanol, and incubated for 1 hour at 37 °C before the addition of LYS-Buffer. The YAC plasmids containing viral cDNA of rSARS-CoV2-E-WT, rSARS-CoV2-E-ΔPBM or rSARS-CoV2-E-MutPBM constructs were digest at the unique restriction site located downstream of the 3ʹ end poly(A) tail using EagI-HF^®^ enzyme (R3505, *New England Biolabs*) following the manufacturer protocol. The cDNA purified by a classical phenol-chloroform process was then transcribed *in vitro* the T7 RiboMax^TM^ Large Scale RNA Production System (P1300, *Promega*). The synthesized RNA was finally purified using a classical phenol-chloroform method, precipitated, and resuspended in Dnase/Rnase free water. 12 μg of complete viral mRNA and 4 μg of pCMV plasmid encoding the viral nucleoprotein (N) gene were electroporated into 8.10^6^ Vero-E6 cells (ATCC #CRL-1586) resuspended in 0.8 mL of Electroporation Solution (MIR50114, *Mirus Bio™ Ingenio™)* using the Gene Pulser Xcell Electroporation System (1652660, *BioRad*) with a pulse of 270V and 950 μF. Cells were then transferred to a T75 culture flask with 12 mL of DMEM supplemented with 2% FCS (v/v) and cultured at 37 °C, 5% CO_2_ for several days until the cytopathic effect (CPE) was observed. The supernatant that corresponds to a P0 stock was harvested, aliquoted and frozen at −80 °C until titration. The viral sequence was controlled by NGS sequencing.

### Viral stock titration

Recombinant viruses’ titer from P0 stock was determined by lysis plaque assay. 1. 10^6^ Vero-E6 cells were seeded to each well of 6-well plates and cultured overnight at 37 °C, 5% CO_2_. Virus was serially diluted in DMEM without FCS and 400 µl diluted viruses were transferred onto the monolayers. The viruses were incubated with the cells at 37 °C with 5% CO_2_ for 1 hour. After the incubation, inoculum was removed, and an overlay medium was added to the infected cells per well. The overlay medium per well contained MEM 1X (prepared from MEM 10X (21430020, *Gibco™*), distilled water (15230162, *Gibco™*), L-glutamine (25030123, *Gibco™*) and gentamicin 10mg/ml (15710064, Gibco™)), 0.25% sodium phosphate (25080094, *Gibco™*), and 1X AVICEL (RC-581*, Dupont*). After a 4 to 6 days of incubation, the plates were stained with crystal violet (V5265, *Merck*) and lyses plaques were counted to assess virus titer.

### Virus growth kinetics

Vero-E6 cells were used to compare the replication kinetics of Wuhan SARS-CoV-2 and rSARS-CoV-2-E-WT viruses with rSARS-CoV2-E-ΔPBM and rSARS-CoV2-E-MutPBM mutant viruses. On the day before infection, 7.10^6^ Vero-E6 cells were seeded onto T75 flask. Cells were washed once with PBS and inoculated with the different viruses in 4 mL of their respective cell culture media for 1 hour at MOI of 0.01, 0.001 or 0.01. Inoculums were then removed, and 12 mL of their respective cell culture media were added in each flask. All cells were maintained at 37 °C in a humidified atmosphere in the presence of 5% CO_2_. Cell-culture supernatants were collected at the indicated time points after infection (8, 24, 32, 48, 56, 72 and 80 hpi) and frozen at −80 °C. Viral titers were obtained by classical TCID_50_ method on Vero-E6 cells after 72 hpi ^82^. Every experiment was performed in triplicate (n=3).

### Maestro Z Impedance experiments

Impedance measurements were conducted using CytoView-Z 96-well electrode plates (*Axion BioSystems*, Atlanta, GA, USA) on a Maestro Z device (*Axion BioSystems*, Atlanta, GA, USA). Prior to cell plating, each well was coated with a 2 mg/mL solution of human fibronectin (F1141, *Merck*) for 1 hour at 37 °C. 100 μL of DMEM containing 5% FCS was added to establish resistance baseline. Vero-E6, at a density of 50 000 cells per well were seeded onto the fibronectin-coated plates in medium supplemented with 5% FCS. The plate was then inserted into the Maestro Z device and resistance at 10 kHz, as a component of impedance, was monitored to observe cell attachment and monolayer stabilization. Infections with recombinant SARS-CoV-2 viruses were performed across a range of MOI from 0.0001 to 1 by incubating cells 1h with 50 μL of inoculum volume. Inoculum was removed and 100 μl of DMEM without FCS were added per well. Resistance kinetics were tracked for up to 96 hpi. Data acquisition and analysis were performed using the Axis Z software.

### Golden Syrian hamster SARS-CoV-2 infection

Male Syrian hamsters (Mesocricetus auratus) of 5–6 weeks of age (average weight 60–80 g) were purchased from Janvier Laboratories and handled under specific pathogen-free. Hamsters were housed by groups of four animals in isolators in a biosafety level-3 facility, with ad libitum access to water and food. Following an acclimation period of one week, animal infections were performed as previously described (PMID: 37495586). Briefly, hamsters were anesthetized with an intraperitoneal injection of ketamine (200 mg/kg, Imalgène 1000, Merial) and xylazine (10 mg/kg, Rompun, Bayer). Infections were performed intranasally by administering 100 µL of physiological solution containing 6.10^4^ PFU of SARS-CoV-2 recombinant viruses. Body weight variation and clinical sign apparition (clinical score) were monitored for four days. At day 3 post-infection (dpi), animals underwent a food finding test to assess olfaction as previously described ^28^. At day 4 post-infection, animals were euthanized with an excess of anesthetics (ketamine and xylazine) and exsanguination ^83^, and samples of nasal turbinates, lungs, olfactory bulbs, cortexes, brain stems and cerebellums were collected and immediately frozen at −80 °C. Lungs, complete head and brain were also collected and fixed in 10% neutral-buffered formalin for histopathological studies.

### SARS-CoV-2 detection in golden hamsters’ tissues

Frozen lungs, nasal turbinates and brain parts (olfactory bulbs, cortexes, brain stems and cerebellums) were weighted and homogenized with 1 mL of DMEM supplemented with 1% penicillin/streptomycin (15140148, *Thermo Fisher Scientific*) in Lysing Matrix M 2 mL tubes (116923050-CF, *MP Biomedicals*) using the FastPrep-24™ system (*MP Biomedicals*). 2 cycles of homogenization at 4.0 m/s during 20 seconds were performed with a resting time of 2 minutes in between. The tubes were centrifuged at 10,000g during 2 minutes at 4 °C, and the supernatants collected. A classical TCID_50_ method on Vero-E6 cells after 72 hpi ^82^ allowed assessment of viral titers. To quantify genomic and sub-genomic Viral RNA loads, RNA was extracted from 125 µL of the supernatants homogenized with 375 µL of Trizol LS (10296028, *Invitrogen*) using the Direct-zol RNA MiniPrep Kit (R2052, *Zymo Research*). Taqman one-step qRT-PCR on E gene (Invitrogen 1 1732-020) was done in a 12.5 μL reaction volume in 384-wells PCR plates using a QuantStudio 6 Flex thermocycler (*Applied Biosystems*). Briefly, 2.5 μL of RNA were added to 10 μL of a master mix containing 6.25 μL of 2X reaction mix, 0.2 µL of MgSO_4_ (50 mM), 0.5 µL of Superscript III RT/Platinum Taq Mix (2 UI/µL) and 3.05 μL of nuclease-free water containing 400 nM of primers and 200 nM of probe. E_sarbeco primers and probe were used to detect genomic RNA (E_Sarbeco_F1 5ʹ-ACAGGTACGTTAATAGTTAATAGCGT-3’; E_Sarbeco_R2 5ʹ-ATATTGCAGCAGTACGCACACA-3’; E_Sarbeco_Probe FAM-5ʹ-ACACTAGCCATCCTTACTGCGCTTCG-3’-TAMRA). For sub-genomic SARS-CoV-2 RNA, detection was achieved by replacing the E_Sarbeco_F1 primer by the CoV2sgLead primer (CoV2sgLead-Fw 5ʹ-CGATCTCTTGTAGATCTGTTCTC-3’). A synthetic gene containing the PCR target sequences was purchased from *Thermo Fisher Scientific*. PCR amplification was performed using Phusion™ High-Fidelity DNA Polymerase (F530, *Thermo Fisher Scientific*) and subsequently transcribed *in vitro* using T7 RiboMax^TM^ Large Scale RNA Production System (P1300, *Promega*). The RNA was quantified using the Qubit RNA HS Assay kit (15958140, *Thermo Fisher Scientific*), normalized, and employed as a standard for determining the absolute copy number of RNA. The amplification process involved an initial incubation at 55 °C for 20 minutes, followed by an initial denaturation step at 95 °C for 3 minutes. This was followed by 50 cycles of denaturation at 95 °C for 15 seconds and annealing/extension at 58 °C for 30 seconds, with a final extension step at 40 °C for 30 seconds. Data was processed using the QuantStudio Design Analysis software.

### Gene expression RT-qPCR

RNA from organs were extracted in trizol using Direct-zol RNA Miniprep kit following the manufacturer protocol (R2050, *Zymo Research*). RNA was reverse transcribed into cDNA using SuperScript™ IV VILO™ Master Mix (11766050, *Invitrogen*). qPCR was performed according to the manufacturer’s protocols for either TaqMan Fast Advanced Master Mix (444457, *Applied Biosystems*) or SYBR Green Master Mix (4309155, *Applied Biosystems*), using 2.5 μL of cDNA (12.5 ng) and golden hamster primer pairs for the genes of interest, as reported in the literature ^28,84^. The qPCR reactions in a final volume of 10 μL were performed in 384-well PCR plates using a QuantStudio 6 Flex thermocycler (*Applied Biosystems*). The amplification conditions were as follows: 95 °C for 20 seconds followed by 45 cycles of 95 °C for 1 second and 60 °C for 20 seconds. Data were treated using the Design and Analysis v.2.8.0 software (*Applied Biosystems*) using *γ-actin g*ene as reference and comparing the n-fold change in expression in the tissues from the infected hamsters with the tissues of the mock-infected group using the 2-ΔΔCt method.

### Histopathology and Immunohistochemistry

Lung fragments, brains and heads fixed 7 days in 10% neutral-buffered formalin were embedded in paraffin. A decalcification treatment was performed on heads consisting of two baths of 4 days within an EDTA based OSTEOMOLL**®** solution (101736, *Merck*). 4 µm thick organ sections were cut and stained with hematoxylin and eosin staining. Lung sections immunohistochemistry was performed on a Bond RX (*Leica^TM^*) using a rabbit polyclonal antibody against SARS Nucleocapsid Protein antibody (1:500, NB100-56576, *Novus Biologicals*) and biotinylated goat anti-rabbit Ig secondary antibody (1:600, E0432, *Dako®, Agilent*). Brain sections immunohistochemistry was performed on a Ventana BenchMark stainer (*Roche^TM^*) using a mouse monoclonal antibody against GFAP (1:500, 6F2, *Dako®, Agilent*), a rabbit polyclonal antibody against fibrinogen (1:500, *Dako®, Agilent*) and a biotinylated secondary antibody included in the detection kit (Ventana DAB Detection Kit 250-001, *Roche^TM^*). Diaminobenzidine (DAB), occasionally combined with alkaline phosphatase (ALP) for double labelling, was used as a chromogen. Slides were then scanned using Axioscan Z1 slide scanner (*Zeiss*), and images were analyzed with the Zen 2.6 software (*Zeiss*).

### MucilAir™ infection

The MucilAir™ model, a pseudostratified bronchiolar epithelium, was purchased from *Epithelix* (Saint-Julien-en-Genevois, France). The model was cultured at 37 °C under a 5% CO_2_ atmosphere with 700 µl of medium in the basal compartment, maintaining the apical side exposed to air to mimic the air-liquid interface. Infections were performed in the apical compartment by incubating the epithelium with 500 000 plaque-forming units of recombinant viruses diluted in medium (EP04MM, *Epithelix*), for 4 hours at 37 °C. After incubation, the viral inoculum was removed, and the epithelium was washed twice with 200 µL of PBS for 5 minutes at 37 °C and once with 200 µL of medium. The apical side was washed every 2-3 days by incubating it with 200 µL of medium for 20 minutes at 37 °C, and the basal medium was replaced every 2–3 days. All supernatants were collected for further analysis.

### Immunofluorescence experiments

MucilAir™ cultures at 4 dpi and A549 ACE2 TMPRSS infected cells were washed twice with PBS and fixed in 4% PFA for 30 minutes. The fixed samples were washed in PBS for 5 minutes three times. MucilAir™ and A549 ACE2 TMPRSS cells were respectively permeabilized during 20 minutes with PBS-TritonX-100 0.5% and PBS-TritonX-100 0.1%. After 3 washes of 5 minutes with PBS-Tween 0.05%, MucilAir™ membranes were incubated in a blocking solution composed by PBS-Tween 0.05%, 1% BSA, 10% normal goat serum (ref) and A549 ACE2 TMPRSS cells were blocked with a solution of PBS-Tween 0.05% with 5% normal goat serum. The permeabilized samples were incubated overnight at 4 °C with primary antibodies diluted in blocking solution: rabbit anti-E 1:250 to 1:500 (HL1443, *Genetex*), mouse anti-GM130 1:250 to 1:500 (610823, *BD Biosciences*), mouse anti-ZO-1 1:50 to 1:100 (610966, *BD Biosciences*), mouse anti-hAfadin 1:100, (MAB78291, *Novus Biologicals*). The next day, after 3 washes of 5 minutes in PBST, samples were incubated 1 hour at 4 °C with conjugated secondary antibody diluted in blocking solution: anti-rabbit antibody conjugated to AF488 1:500 to 1:1000 (A11035, *Invitrogen*) and anti-mouse antibody conjugated to AF594 1:500 to 1:1000 (A21125, *Invitrogen*). After 3 washes of 5 minutes, samples were mounted in Fluoroshield™ with DAPI (F6057, *Merck*). Stainings were observed with a LSM700 confocal microscope (*Zeiss*). Acquired images were processed using ZEN lite software (*Zeiss*).

### Pull Down assays

Subconfluent cultures of HEK 293 cells were transiently transfected with a plasmid encoding GFP or GFP-tagged ZO-1 full-length protein using the calcium phosphate method. In parallel, subconfluent monolayer cultures of Vero-E6 cells were infected with rSARS-CoV2-E-WT or rSARS-CoV2-E-MutPBM viruses for 48 hours at a MOI of 0.01. Both cell cultures were then scraped, lysed using RIPA Lysis Buffer (sc-24948, *Santa Cruz*), and centrifuged at 13,000 rpm for 10 minutes at 4 °C to pellet cell debris. Soluble detergent extracts from the GFP constructs expressing cells were incubated with GFP resins for 2 hours at 4 °C with gentle shaking, and then washed 3 times with a washing buffer containing PBS supplemented with 200 mM NaCl and 0.1% Triton X-100. The GFP construct coated beads were then incubated with mock or infected cell lysates during 2 hours at 4 °C, followed by three washes. The eluted proteins were processed for Western blot analysis using GFP antibody (NB600-313, *Novus Biologicals*) and envelope protein antibody (HL1443, *Genetex*).

### RNA libraries and sequencing

Libraries were built using a Illumina Stranded mRNA library Preparation Kit *(Illumina*, USA) following the manufacturer’s protocol from the RNA extractions of brainstem and lung samples from non-infected hamsters or infected with the recombinant viruses (E-WT, E-ΔPBM or E-MutPBM), producing 4 replicates per group. Quality control was performed on an BioAnalyzer (*Agilent*). Sequencing was performed on one NovaSeq X lane 10B300 (*Illumina*) to obtain 150 base paired end reads. Unfortunately, the libraries for the 4 replicates of the E-MutPBM infected lung failed and had to be performed a second time and sequenced in another NextSeq 2000 sequencing run to obtain 100 base single-end reads.

### RNA-seq analysis

The RNA-seq analysis was performed with Sequana 0.16.11 ^85^. We used the RNA-seq pipeline 0.19.2 (https://github.com/sequana/sequana_rnaseq) built on top of Snakemake 7.32.4 ^86^. Briefly, reads were trimmed from adapters using Fastp 0.22.0 ^87^ then mapped to the golden hamster MesAur1.0.100 genome assembly from Ensembl using STAR 2.7.10a ^88^. FeatureCounts 2.0.1 ^89^ was used to produce the count matrix, assigning reads to features using corresponding annotation from Ensembl with strand-specificity information. Quality control statistics were summarized using MultiQC 1.17 ^90^. Statistical analysis on the count matrix was performed to identify differentially regulated genes comparing each recombinant viruses infected tissues versus the mock group and including a batch effect to the model corresponding to the replicates. Clustering of transcriptomic profiles were assessed using a Principal Component Analysis (PCA). Differential expression testing was conducted using DESeq2 library 1.34.0 ^91^ scripts indicating the significance (Benjamini-Hochberg adjusted p-values, false discovery rate FDR < 0.05) and the effect size (fold-change) for each comparison. Finally, enrichment analysis from the differentially regulated gene lists was performed using modules from Sequana. The KEGG pathways enrichment uses GSEApy 1.1.1 ^92^, EnrichR ^93^ and KEGG database ^94^. Programmatic accesses to online web services were performed via BioServices 1.11.2 ^95^.

### Statistics and reproducibility

Statistical analysis was performed using Prism 10 (*GraphPad*, version 10.1.1, San Diego, USA). Quantitative data was compared across groups using Kruskal-Wallis’ test followed by Dunn’s multiple comparisons test (horizontal lines indicate median with the interquartile range). Statistical significance was assigned when *p* values were <0.05 and is indicated in the figures. Randomization and blinding were not possible for *in vivo* experiments due to pre-defined housing conditions (separated isolators between animals infected by each virus strains). *Ex vivo* and *in vitro* analyses were blinded (coded samples). The nature of statistical tests and the number of experiments or animals (n) are reported in the figure legends.

## Supporting information

all supplementary material

## Acknowledgements

This work was supported by: Institut Pasteur TASK FORCE COVID19, Institut Pasteur’s Programme Fédérateur de Recherche 5 (PFR5-Functional Genomics of the Viral Cycle funded by Fondation d’entreprise Michelin donation) and Institut Pasteur EID Junior Call. Flavio Alvarez is recipient of a fellowship from the Institut Pasteur’s Programme Fédérateur de Recherche 5 (PFR-5 - Functional Genomics of the Viral Cycle). The production of recombinant viruses was performed thanks to the techniques developed by the Institut Pasteur’s Programme Fédérateur de Recherche 1 (PFR-1 - Reverse Genetics). We would like to acknowledge Caroline Demeret for the fruitful discussions about SARS-CoV-2 protein interactome. We thank Remy Robinot and Lisa Chakrabarti (Virus and Immunity Unit, Institut Pasteur) for their advices on the Lung-on-chip model. We would like to thank Axion Biosystem for giving us the opportunity to experiment with the Maestro machine, and especially Emma David and Nicolas Roy for their support and assistance. Part of this work was performed at the Histological core facility, UtechS Photonic BioImaging (PBI) and Biomics platforms of the Institut Pasteur. We acknowledge the work performed in transcriptomic experiments by Elodie Turc and Iakov Vintrenko, members of Biomics Platform directed by Marc Monot (C2RT, Institut Pasteur) supported by France Génomique (ANR-10-INBS-09) and IBISA. Figure 2A was generated using Biorender software under Institut Pasteur license.

