## Supplementary material for "The SARS-CoV-2 envelope PDZ Binding Motif acts as a virulence factor disrupting host’s epithelial cell-cell junctions": all supplementary material

**⥉These authors contributed equally**

**# These authors contributed equally**

| Rescue P0 rSARS-COV-2-E-ΔPBM | | | |
|---|---|---|---|
| Protein | Nucleotide | Amino acid | Frequency in the viral population (%) |
| ORF1 a-b | A6524G | T2087A | 11.6 |
| Envelope | Δ26458-26469 | ΔDLLV 72-75 | 100 |
| | Δ26458-26469 ; C26456T ; T26457C | P71L ; ΔDLLV 72-75 | 3 |

| Rescue P0 rSARS-COV-2-E-MutPBM | | | |
|---|---|---|---|
| Protein | Nucleotide | Amino acid | Frequency in the viral population (%) |
| Envelope | A26459G ; C26461G ; T26462G ; C26464G ; T26465G ; G26466A ; T26468G ; C26469T | DLLV → GGGG 72-75 | 100 |

**Supplementary Table 1. Sequencing analysis of the SARS-CoV-2 recombinant viruses**. Next Generation Sequencing (NGS) was performed in P0 rescue of rSARS-CoV-2-E-ΔPBM and rSARS-CoV-2-E-MutPBM virus. Mutations of the envelope protein and elsewhere in the viral genome above a frequency of 10% were considered.

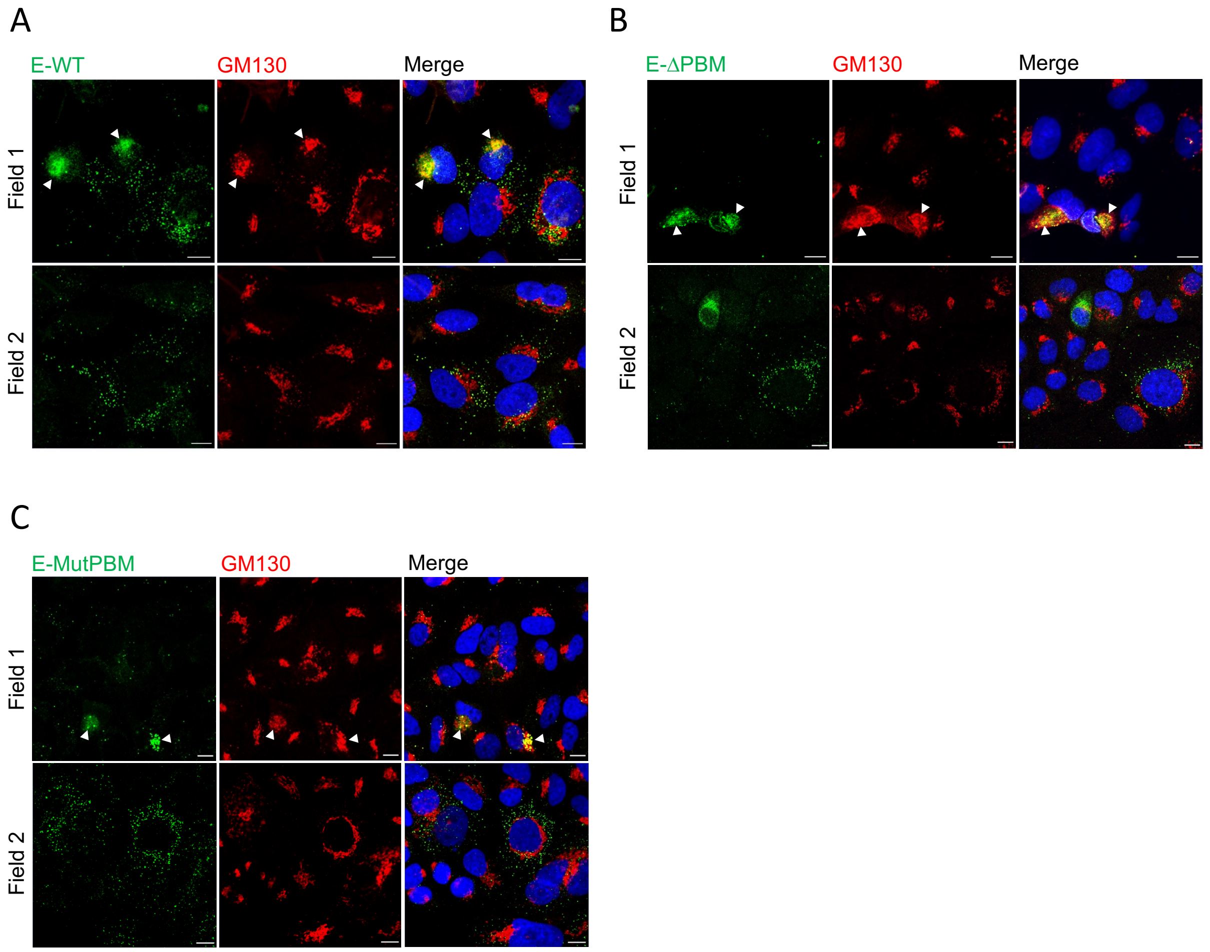


**Supplementary Figure 1. Subcellular localization of the Envelope protein during SARS-CoV-2 infection in lung epithelial cell model.** Visualisation of E protein in A549 ACE2 TMPRSS cell by immunofluorescence assay. Staining was performed on 48h infected cells at a 0.1 MOI with recombinant E-WT (**A**), E-ΔPBM (**B**) or E-MutPBM viruses (**C**). Fields 1 images of E protein are related to Golgi location whereas Fields 2 images exhibit cytoplasmic vesicular distribution of E (scale bar = 10 µm). Hoechst: nuclei (blue); GM130: golgi apparatus (red); E-WT/E-ΔPBM/E-MutPBM: recombinant SARS-CoV-2 envelope protein (green). Arrowheads indicate E-golgi colocalizations. Related to Figure 2

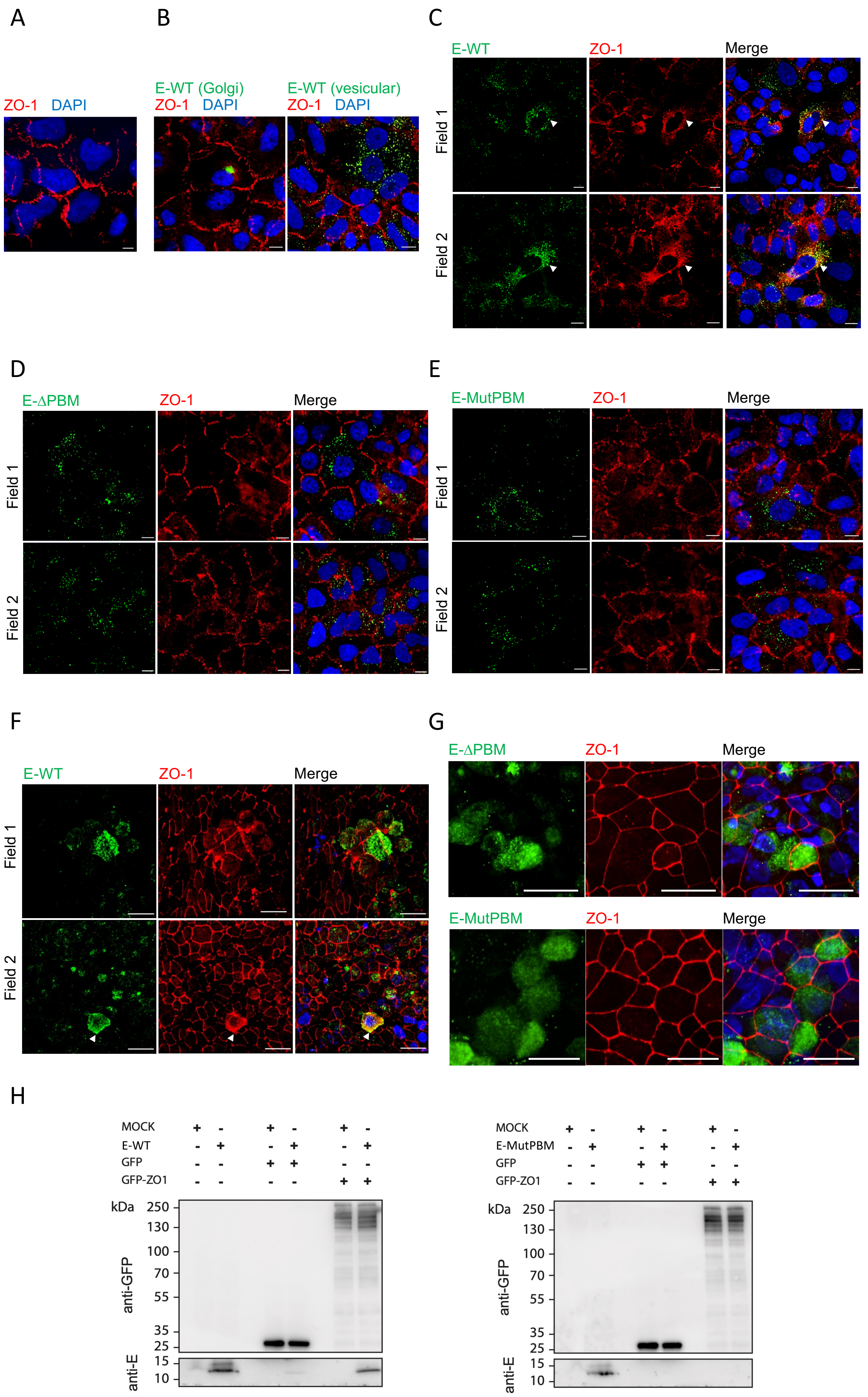

A
B
C
ZO-1 DAPI
E-WT (Golgi)
ZO-1 DAPI
E-WT (vesicular)
ZO-1 DAPI
E-WT
ZO-1
Merge
Field 1
Field 2
D
E
E-ΔPBM
ZO-1
Merge
E-MutPBM
ZO-1
Merge
Field 1
Field 2
F
G
E-WT
ZO-1
Merge
E-ΔPBM
ZO-1
Merge
E-MutPBM
ZO-1
Merge
H
MOCK + - + - + -
E-WT - + - + - +
GFP - - + + - -
GFP-ZO1 - - - - + +
kDa 250 130 100 70 55 35 25
anti-GFP
15 10
anti-E
MOCK + - + - + -
E-MutPBM - + - + - +
GFP - - + + - -
GFP-ZO1 - - - - + +
kDa 250 130 100 70 55 35 25
anti-GFP
15 10
anti-E

**Supplementary Figure 2. Subversion of ZO-1 protein during SARS-CoV-2 infection in epithelial cell line in a PBM dependent manner. A.B.C.** Visualization by immunofluorescence of ZO-1 protein disruption in A549 ACE2 TMPRSS2 cells in correlation with E protein localization. Staining was performed on cells that were either non-infected **(A)** or infected for 48 hours at a MOI of 0.1 with the recombinant E-WT virus **(B)**. Arrowheads indicate regions of colocalization between E protein and ZO-1 **(C)**. **D.E.** Observation by immunofluorescence of a significantly reduced disruption of the ZO-1 network in A549 ACE2 TMPRSS2 cells infected for 48 hours with either the E-ΔPBM **(D)** or E-MutPBM **(E)** recombinant viruses. Hoechst: nuclei (blue); ZO-1: zonula occludens 1 protein (red); E-WT/E-ΔPBM/E-MutPBM: recombinant SARS-CoV-2 viruses envelope protein (green) (scale bar = 10 µm). **F.G.** Supplementary visualization of the ZO-1 protein network in correlation with E protein localization within the MucilAir model using immunofluorescence staining. Staining was performed on epithelium fixed at 4 days post-infection (dpi) either with 250,000 pfu of E-WT virus **(F)** or PBM lacking recombinant viruses (E-ΔPBM and E-MutPBM) **(G)**. Images are reconstituted in 3D from Z stack acquisitions (scale bar = 20 µm). Hoechst: nuclei (blue); ZO-1: zonula occludens 1 (red); Envelope: recombinant SARS-CoV-2 envelope protein (green). **H.** Western blot of GFP pull-down. GFP or GFP-ZO-1 from transfected HEK293 cell lysates were immobilised on GFPtrap resin and incubated with Vero-E6 cell lysates non infected or infected 48h at a 0.01 MOI with recombinant E-WT (left panel) or E-MutPBM viruses (right panel). The bound fractions were analysed by immunoblotting using anti-GFP (top panel) and anti-E (bottom panel) antibodies. Related to Figure 2

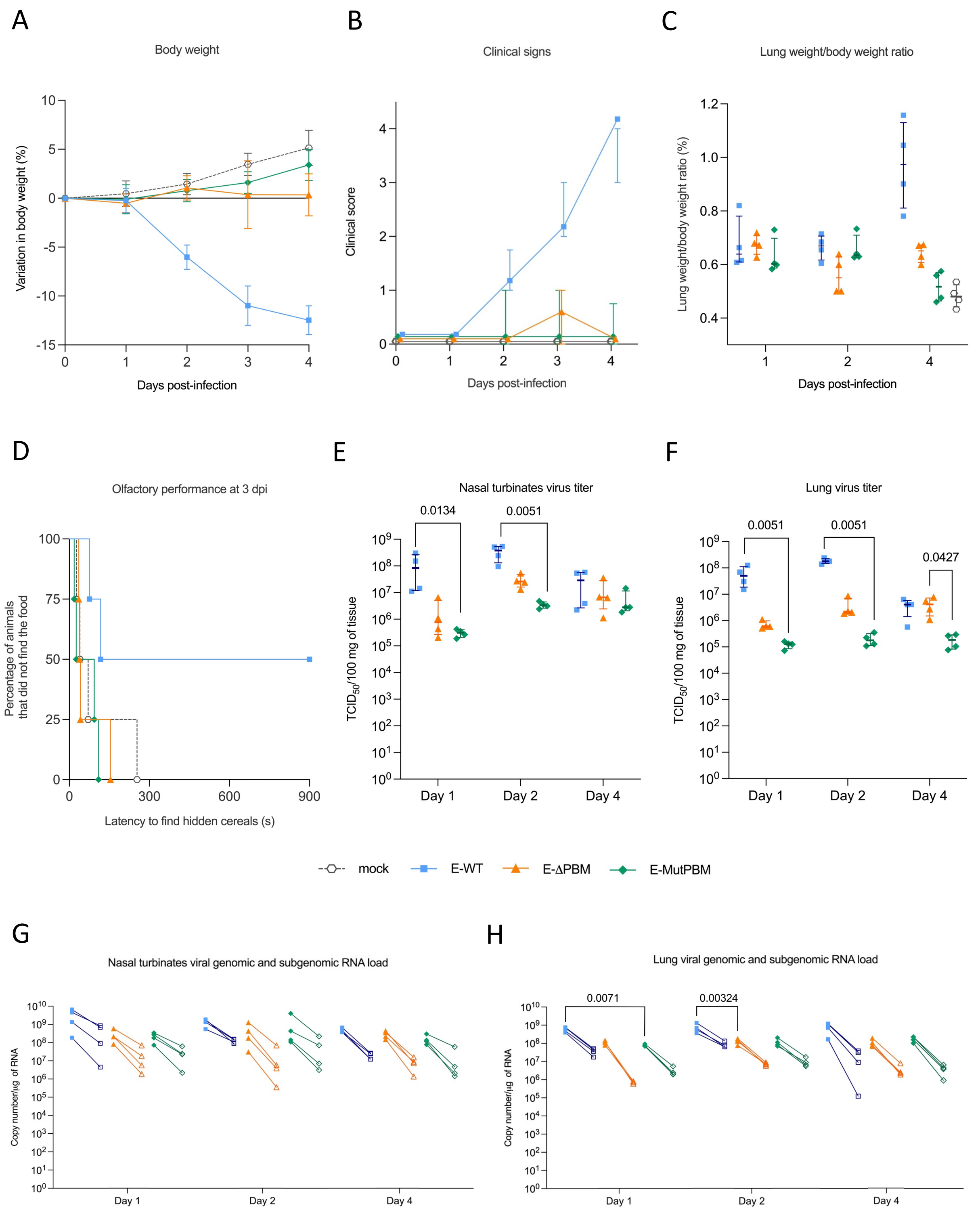


**Supplementary Figure 3 : Kinetics of clinical profile and viral metrics of hamsters infected with recombinant wild-type SARS-CoV-2 (E-WT) and PBM lacking viruses (E-ΔPBM and E-MutPBM). A.** Body weight variation over the four days post-infection. **B.** Clinical score over the four days post-infection. The clinical score is based on a cumulative 0–4 scale: ruffled fur; slow movements; apathy; and absence of exploration activity. **C.** Lung weight-to-body weight ratio at 1, 2 and 4 dpi. Horizontal lines indicate median with the interquartile range (n=4/group). **D.** Olfactory performance measured at 3 days post-infection (dpi). The olfaction test is based on the hidden (buried) food finding test. Curves represent the olfactory performance of animals during the test (n= 4/group). **E.F.** Infectious viral titers in nasal turbinates **(E)** and lung **(F)** at 1,2 and 4 dpi expressed as $TCID_{50}$ per 100 mg of tissue. Horizontal lines indicate median and the interquartile range (n=4/group). Kruskal-Wallis test followed by the Dunn's multiple comparisons test (the adjusted p value is indicated when significant). **G.H.** Viral genomic and subgenomic RNA load detected in nasal turbinates **(G)** and lung **(H)** at 1,2 and 4 dpi. Horizontal lines indicate median and the interquartile range. Lines connect symbols from the same animals (n=4/group). Kruskal-Wallis test followed by the Dunn's multiple comparisons test (the adjusted p value is indicated when significant). (n=4/group). Related to Figure 3

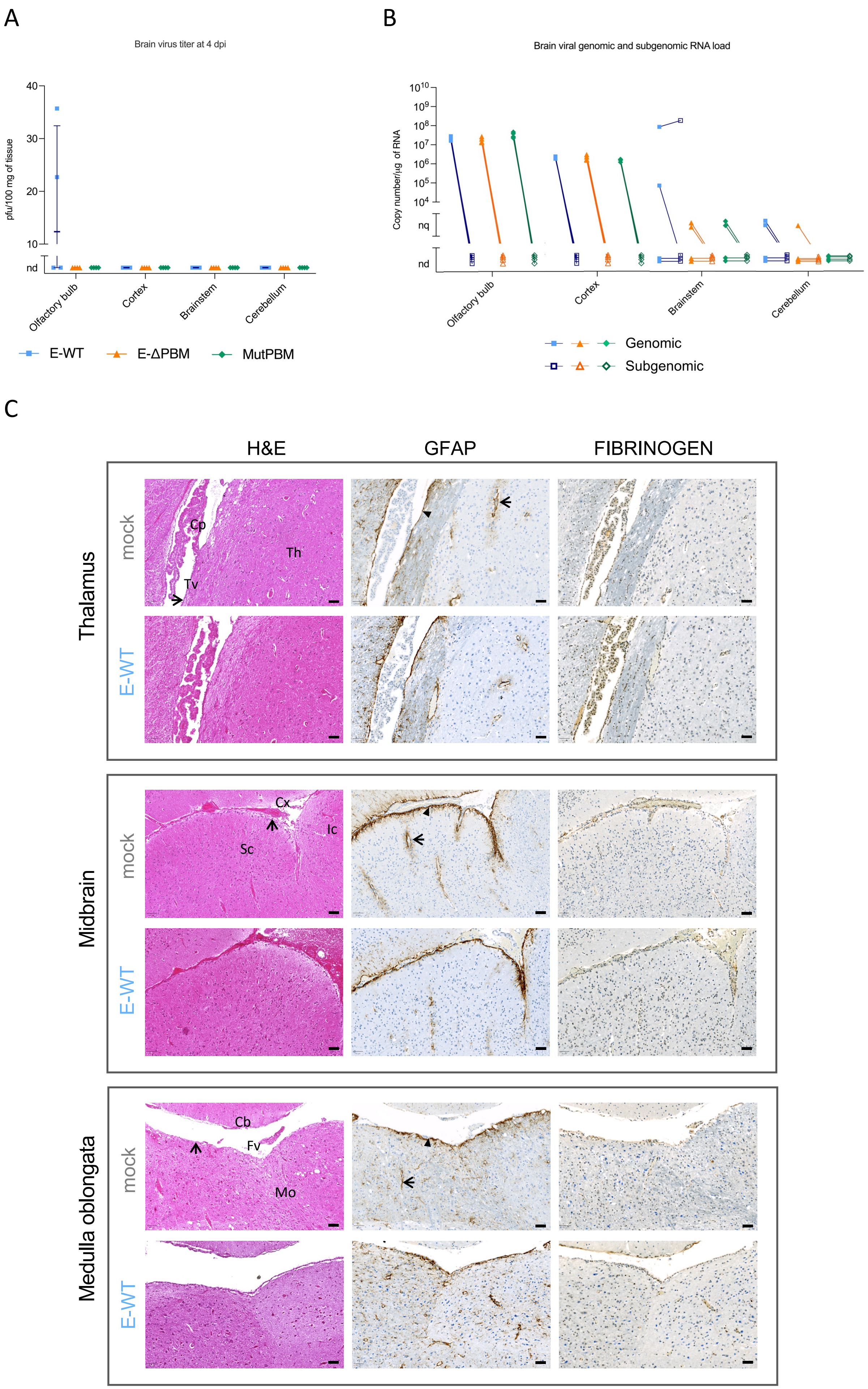
A
Brain virus titer at 4 dpi
pfu/100 mg of tissue
40
30
20
10
nd
Olfactory bulb
Cortex
Brainstem
Cerebellum
E-WT
E-ΔPBM
MutPBM
B
Brain viral genomic and subgenomic RNA load
Copy number/µg of RNA
$10^{10}$
$10^{9}$
$10^{8}$
$10^{7}$
$10^{6}$
$10^{5}$
$10^{4}$
nq
nd
Olfactory bulb
Cortex
Brainstem
Cerebellum
Genomic
Subgenomic
C
H&E
GFAP
FIBRINOGEN
Thalamus
mock
E-WT
Cp
Th
Tv
Midbrain
mock
E-WT
Cx
Ic
Sc
Medulla oblongata
mock
E-WT
Cb
Fv
Mo

**Supplementary Figure 4. Viral quantification and brain histopathology in hamsters infected with SARS-CoV-2 recombinant virus (E-WT) at 4 dpi (days post-infection) A.** Infectious viral titers in brain regions (olfactory bulb, cortex, brainstem and cerebellum) at 4 day post-infection (dpi) expressed as pfu per 100 mg of tissue. Horizontal lines indicate median and the interquartile range (n=4/group). Kruskal-Wallis test followed by the Dunn's multiple comparisons test (the adjusted p value is indicated when significant). **B.** Viral genomic and subgenomic RNA load detected in brain regions at 4 dpi. Genomic and subgenomic viral RNA were assessed based on the E gene sequence. Horizontal lines indicate median and the interquartile range. Lines connect symbols from the same animals (n=4/group). Kruskal-Wallis test followed by the Dunn's multiple comparisons test (the adjusted p value is indicated when significant). (n=4/group, nd: not detected, nq: not quantifiable, detected outside the standard curve). **C.** Histopathology and immunohistochemical analysis of brain tissue from non-infected hamsters and those infected with WT SARS-CoV-2 recombinant virus (E-WT) at 4 dpi. Representative images include Hematoxylin and Eosin (H&E) staining (left panels), Glial fibrillary acidic protein (GFAP) immunohistochemistry (middle panels), and FIBRINOGEN staining (right panels) in the thalamus (top row), midbrain (middle row), and medulla oblongata (bottom row) regions of the brain. Scale bar = 50 µm; n = 4 per group. Abbreviations: Cp: choroid plexus; Tv: third ventricle; Th: thalamus; Cx: cerebral cortex; Sc: superior colliculus; Ic: inferior colliculus; Cb: cerebellum; Fv: fourth ventricle; Mo: medulla oblongata. In the H&E images, arrows indicate the ependymal layer in the thalamus, leptomeninges in the midbrain, and ependymal layer in the medulla oblongata. In the immunohistochemistry images, arrows highlight perivascular astrocyte foot processes, while arrowheads denote subependymal astrocyte foot processes.

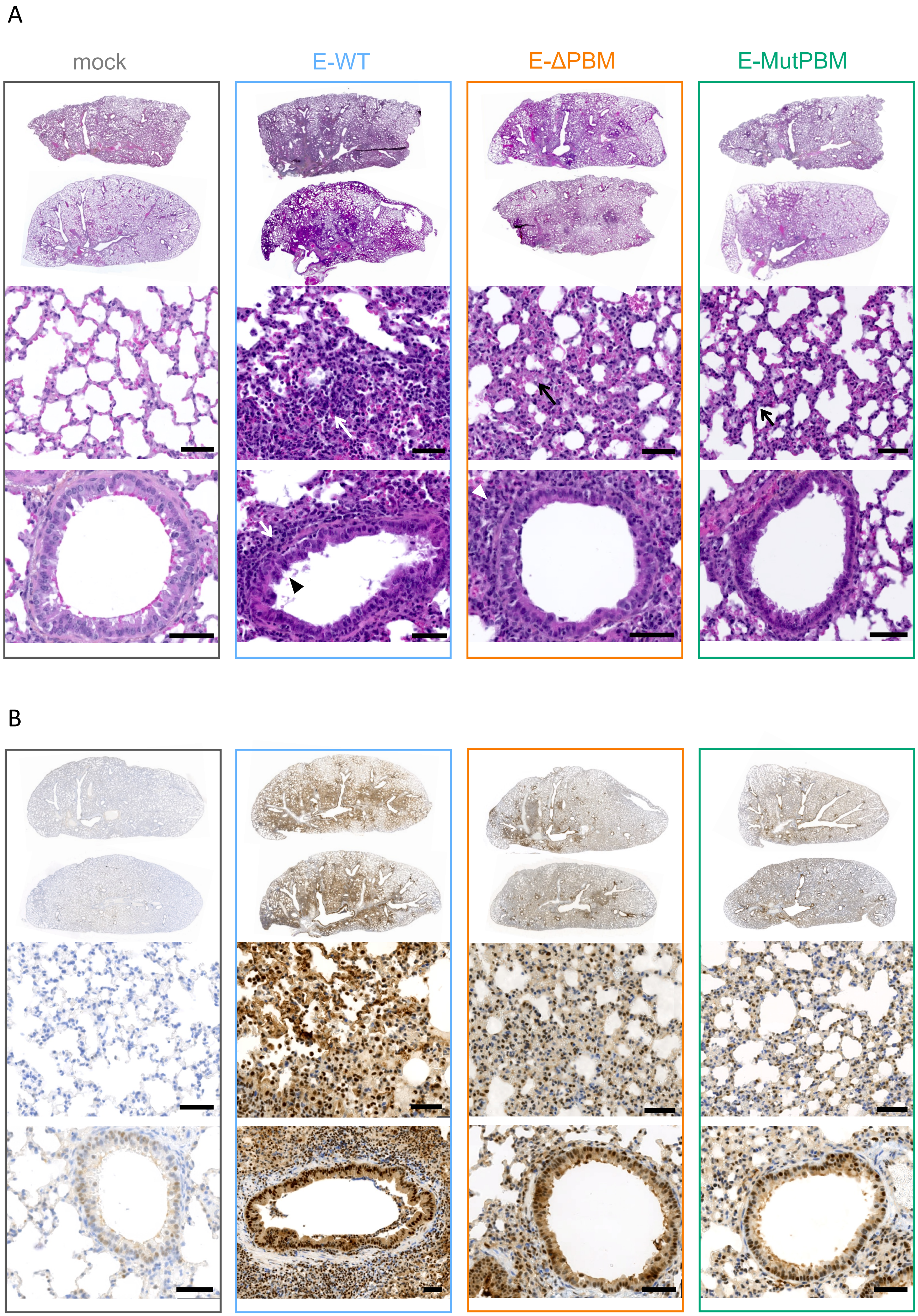
A
mock
E-WT
E-ΔPBM
E-MutPBM
B

**Supplementary Figure 5. Histopathology and immunohistochemical study of lungs from hamsters infected with WT SARS-CoV-2 recombinant virus (E-WT) and PBM lacking viruses (E-ΔPBM and E-MutPBM) at 4 dpi. A.** Representative images of Hematoxylin and Eosin (H&E) stained-whole lung sections (upper panels), alveoli (middle panels) and bronchiolar epithelium (bottom panels). Arrows indicate inflammation and arrowhead correspond to epithelial damages. **B.** Representative images of whole lung sections (upper panels), alveoli (middle panels) and bronchiolar epithelium (bottom panels) immuno-stained with SARS-CoV-2 Nucleocapsid antibody. Scale bar are indicated under images (n=4/group). Related to Figure 3

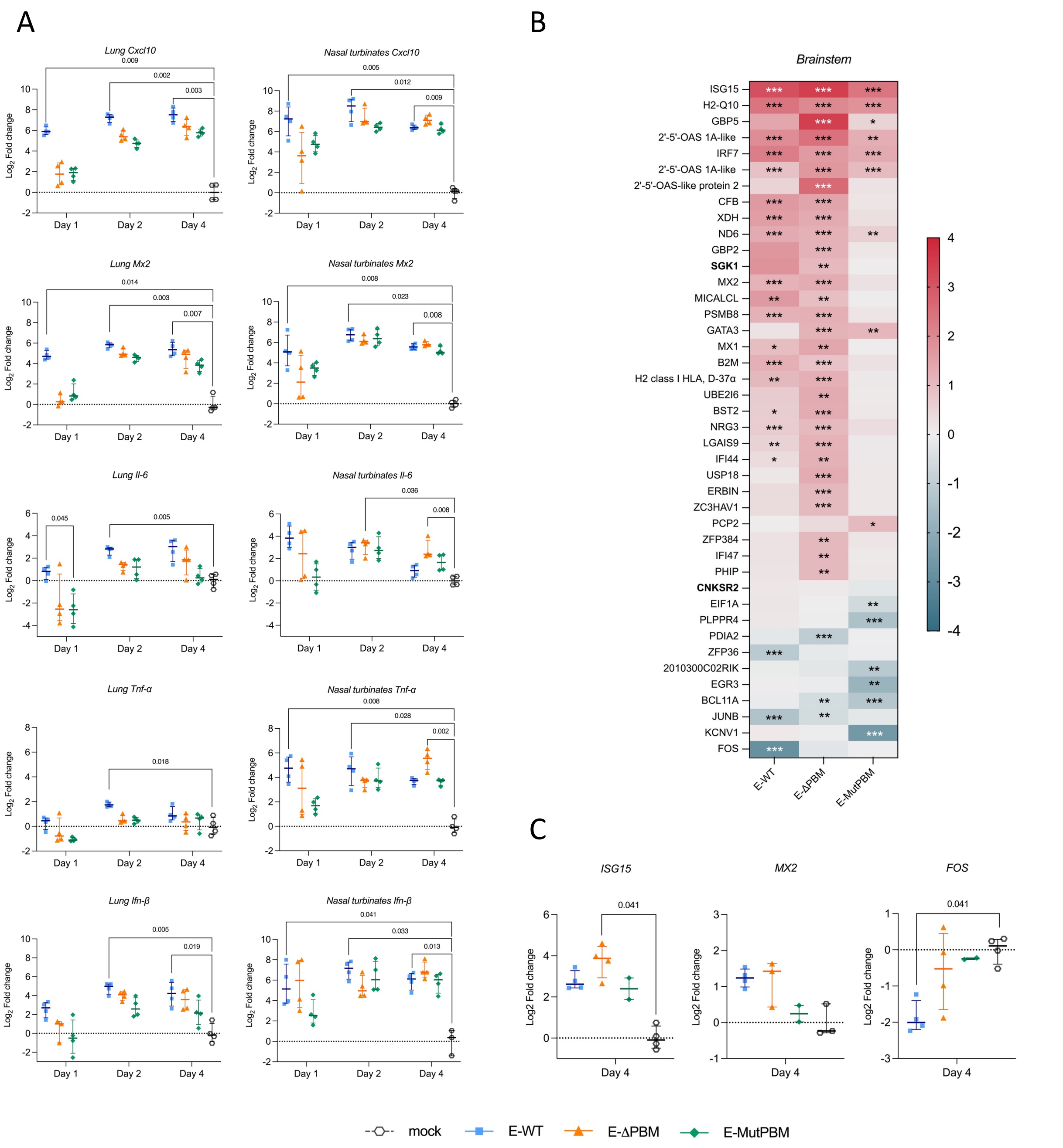


**Supplementary Figure 6. Pro-inflammatory and Differential Gene Expression in Hamster Tissues Following Infection with Recombinant Viruses. A.** Validation of pro-inflammatory gene targets, presented as $Log_2$ fold change in expression, in the lungs (left panels) and nasal turbinates (right panels) of hamsters infected with E-WT, E-ΔPBM, and E-MutPBM, relative to mock-infected controls. Expression levels of Cxcl10, Mx2, Il-6, Ifn-β, and Tnf-α were quantified at 1, 2, and 4 days post-infection (dpi). Horizontal lines indicate the median and interquartile range (n = 4 per group). Statistical significance was assessed using the Kruskal-Wallis test followed by Dunn's multiple comparisons test, with significant p-values indicated ($p<0.05$). **B.** Heatmap displaying all differentially expressed genes in the brainstem of hamsters infected with E-WT, E-ΔPBM, and E-MutPBM, compared to mock-infected controls at 4 days post-infection (dpi), with an absolute fold change > 2. Asterisks (*) correspond to genes with a Benjamini–Hochberg-adjusted p-value < 0.05 in comparisons between each recombinant virus and the mock-infected group. The color gradient represents the $log_2$ fold change in gene expression between infected and mock-infected animals. **C.** Validation of gene targets, presented as $Log_2$ fold change in expression, in the brainstem of hamsters infected with E-WT, E-ΔPBM, and E-MutPBM, relative to mock-infected controls. Expression levels of ISG15, Mx2 and Fos were quantified at 4 days post-infection (dpi). Horizontal lines indicate the median and interquartile range. Statistical significance was assessed using the Kruskal-Wallis test followed by Dunn's multiple comparisons test, with significant p-values indicated ($p<0.05$). Related to Figure 4.
